# The neurobiology of agrammatic sentence comprehension: a lesion study

**DOI:** 10.1101/192245

**Authors:** Corianne Rogalsky, Arianna N. LaCroix, Kuan-Hua Chen, Steven W. Anderson, Hanna Damasio, Tracy Love, Gregory Hickok

**Author notes:** Corresponding author: Corianne Rogalsky, Department of Speech and Hearing Science, Arizona State University, P.O. Box 870102, Tempe, AZ 85287-0102.

## Abstract

Broca’s area has long been implicated in sentence comprehension. Damage to this region is thought to be the central source of “agrammatic comprehension” in which performance is substantially worse (and near chance) on sentences with noncanonical word orders compared to canonical word order sentences (in English). This claim is supported by functional neuroimaging studies demonstrating greater activation in Broca’s area for noncanonical versus canonical sentences. However, functional neuroimaging studies also have frequently implicated the anterior temporal lobe (ATL) in sentence processing more broadly, and recent lesion-symptom mapping studies have implicated the ATL and mid temporal regions in agrammatic comprehension. The present study investigates these seemingly conflicting findings in 66 left hemisphere patients with chronic focal cerebral damage. Patients completed two sentence comprehension measures, sentence-picture matching and plausibility judgments. Patients with damage including Broca’s area (but excluding the temporal lobe; n=11) on average did not exhibit the expected agrammatic comprehension pattern, e.g. their performance was > 80% on noncanonical sentences in the sentence-picture matching task. Patients with ATL damage (n=18) also did not exhibit an agrammatic comprehension pattern. Across our entire patient sample, the lesions of patients with agrammatic comprehension patterns in either task had maximal overlap in posterior superior temporal and inferior parietal regions. Using voxel-based lesion symptom mapping (VLSM), we find that lower performances on canonical and noncanonical sentences in each task are both associated with damage to a large left superior temporal-inferior parietal network including portions of the ATL, but not Broca’s area. Notably however, response bias in plausibility judgments was significantly associated with damage to inferior frontal cortex, including gray and white matter in Broca’s area, suggesting that the contribution of Broca’s area to sentence comprehension may be related to task-related cognitive demands.

## Introduction

At the turn of the 20^th^ century a then decades-long debate persisted regarding the functional role of Broca’s area in speech. Broca had proposed in the 1860s that the “foot of the third frontal convolution”—what we call Broca’s area today—was the seat of articulate speech. However, in 1905 Pierre Marie authored a famous paper declaring that “The third frontal convolution plays no special role in the function of language” (Marie, 1906), presenting evidence that Broca’s area could be damaged with no effect on articulate speech and conversely that deficits in articulate speech could be documented without damage to Broca’s area. Modern data has at the very least confirmed the confusion, if not resolved the debate fully. Damage to Broca’s area *alone* does not cause *chronic* Broca’s aphasia nor significant chronic disruption of speech articulation (Mohr et al. 1978; Mohr, 1976) and Broca’s aphasia can be caused by lesions sparing Broca’s area (Fridriksson, Bonilha & Rorden, 2007).

Approximately a century after Broca’s initial claims regarding his namesake brain area, a new function was tied to the region after it was discovered that Broca’s aphasics are impaired at comprehending sentences that require subtle syntactic analysis, such as semantically reversible sentences with non-canonical word order (*It was the cat that the dog chased* or *The dog was chased by the cat*) compared to more typical subject-verb-object orderings in English (Bradley, Garrett & Zurif, 1980; Caramazza and Zurif, 1976). Notwithstanding the fact that Broca’s aphasia holds a tenuous relation to Broca’s area and the fact that people with conduction aphasia (and therefore posterior temporal-parietal lesions) have the very same sentence comprehension trouble (Caramazza and Zurif, 1976), Broca’s area became the theoretical centerpiece in models of the neurology of syntax generally (Zurif, 1980) and the basis of aspects of sentence comprehension in particular (Caplan & Futter, 1986; Grodzinsky, 1986). But like a century ago, the battle over Broca’s area” (Grodzinsky & Santi, 2008), continues to rage, this time over its role in sentence comprehension. Early dissenters attacked the specificity of syntactic function, showing that Broca’s aphasics could judge the grammaticality of sentences quite well (Linebarger, Schwartz & Saffran, 1983; Wulfeck, 1988). This led to more refined theories regarding which syntactic functions are affected (Grodzinsky & Santi, 2008; Grodzinsky, 1986, 2000).

In the last two decades, the debate has been fueled by evidence from functional imaging. Some groups have shown what appears to be rather specific involvement of Broca’s area during aspects of sentence comprehension (Caplan, Alpert & Waters, 1998, 1999; Caplan, Alpert, Waters & Olivieri, 2000; Dapretto & Bookheimer, 1999; Just, Carpenter, Keller, Eddy & Thulborn, 1996; Stromswold, Caplan, Alpert & Rauch, 1996; Grodzinsky & Santi, 2008; Santi & Grodzinsky, 2007a,b; Fedorenko, Behr & Kanwisher, 2011; Fedorenko & Kanwisher, 2011; Rogalsky, Almeida, Sprouse & Hickok, 2015) while others point to results arguing that the effects are confounded with working memory load differences (Just et al. 1996; Kaan & Swaab, 2002; Rogalsky, Matchin & Hickok, 2008; Rogalsky & Hickok, 2011) or cognitive control demands (Novick, Trueswell & Thompson-Schill, 2005). Still other studies comparing activations in response to sentences versus non-sentence controls, such as word lists or unintelligible acoustically-matched speech, have not reported Broca’s area involvement consistently (Mazoyer et al. 1993; Stowe et al. 1998; Crinion, Lambon-Ralph, Warburton, Howard & Wise, 2003; Humphries, Love, Swinney & Hickok, 2005; Humphries, Binder, Medler & Liebenthal, 2006; Friederici, Meyer & von Cramon, 2000; Friederici, Kotz, Scott & Obleser, 2010; Spitsyna, Warren, Scott, Turkheimer & Wise, 2006; Scott, Blank, Rosen & Wise, 2000; Rogalsky & Hickok, 2009; Rogalsky, Rong, Saberi & Hickok 2011; Awad, Warren, Scott, Turkheimer & Wise, 2007).

Unlike Broca’s area, the anterior temporal lobe (ATL) (Mazoyer et al. 1993; Stowe et al. 1998; Crinion et al. 2003; Humphries et al. 2005, 2006; Friederici et al. 2000, 2010; Spitsyna et al. 2006; Scott, Blank, Rosen & Wise, 2000; Rogalsky & Hickok, 2009; Rogalsky et al. 2011, 2015; Awad et al. 2007) as well as, in some studies, portions of the posterior temporal lobe and/or temporal-parietal junction (Vandenberghe, Nobre & Price, 2002; Humphries et al. 2005, 2006; Humphries, Binder, Medler & Liebenthal, 2007; Snijders et al. 2009; Friederici, 2011), have been shown to be sensitive to sentences compared to a variety of non-sentence control conditions. Portions of the ATL (mostly in the temporal pole, i.e. Brodmann area 38), are widely considered to be a multi-modal semantic hub (Simmons & Martin, 2009; Bonner & Price, 2013); thus, perhaps the ATL’s contribution to sentence comprehension may be due to lexical-semantic processing. However, there is strong neuroimaging evidence suggesting that the ATL’s contributions to sentence comprehension are not solely lexical in nature: portions of the ATL respond more to pseudoword sentences than to pseudoword lists (Humphries et al. 2006; Rogalsky et al. 2011), more to syntactic errors than semantic errors (Friederici, Ruschmeyer, Hahne & Fiebach, 2003; Herrmann, Obleser, Kalberlah, Haynes & Friederici, 2012), and more to idioms than literal sentences (Lauro, Tettamanti, Cappa & Papagno, 2008). The ATL also is more activated by sentence prosody than list prosody (Humphries et al. 2005). However, Broca’s area, but not the ATL, has been found to be more responsive to noncanonical sentences compared to canonical sentences (Caplan et al. 1998, Stromswold et al. 1996; Rogalsky et al. 2008; Rogalsky & Hickok 2011; Ben-Shachar, Palti & Grodzinsky, 2004; see Santi & Grodzinsky (2012) for one notable exception).

Large-scale lesion mapping methods developed in the 2000s (Bates et al. 2003) have offered some hope of resolving the murky state of affairs regarding the neural basis of sentence comprehension. The few such studies that have been undertaken so far mainly implicate temporal-parietal regions in sentence comprehension (Dronkers, Wilkins, van Valin, Redfern & Jaeger, 2004; Thothathiri, Kimberg & Schwartz, 2012; Magnusdottir et al. 2013; Pillay, Binder, Humphries, Gross & Book, 2017). For example, Dronkers, et al. (2004) investigated auditory sentence comprehension using subtests of the CYCLE-R clinical battery, in 64 English-speaking chronic stroke patients with left hemisphere damage. Their recruitment was not based on aphasia diagnosis; their sample contained both patients with and without aphasia. Their voxel-based lesion symptom mapping (VLSM) results implicated both the ATL and Broca’s area in the comprehension of sentences, but noncanonical and canonical sentences were not examined separately or compared, so the relative contributions of Broca’s area and the ATL to noncanonical and canonical sentence comprehension remains unclear.

A subsequent VLSM study (Magnusdottir et al. 2013) did examine noncanonical versus canonical sentence comprehension in 50 Icelandic-speaking patients with acute left hemisphere strokes; most testing was completed within three days of admission to the hospital. Using a sentence-picture matching task, Magnusdottir et al. measured performance on sentences with noncanonical word order as well as on sentences with canonical word order. They report that overall sentence comprehension implicates a large temporal-parietal-occipital area but not any frontal regions including Broca’s area. Their analysis within each sentence type found that canonical sentences implicated posterior temporal and inferior parietal areas, while noncanonical sentences implicated Broca’s area as well as more anterior and inferior temporal lobe areas. They then computed a VLSM of a direct contrast of canonical > noncanonical performance, i.e. an agrammatic comprehension pattern, which identified an ATL region in which damage is associated with impaired performance on noncanonical sentences compared to canonical sentences. The authors conclude that the ATL “plays a crucial role in syntactic processing,” and that Broca’s area contributes as well.

Thothathiri et al. (2012) also examined canonical and noncanonical sentence comprehension in 79 chronic stroke patients with aphasia. Their VLSM analyses identified posterior temporal-inferior parietal regions as critical for both canonical and noncanonical sentences. Thothathiri et al. also conducted a VLSM of the difference between canonical and noncanonical scores (i.e. canonical > noncanonical performance) but found no voxels that reached significance. A region-of-interest follow up, however, implicated posterior temporo-parietal regions. Broca’s area was not implicated in canonical, noncanonical, or canonical > noncanonical performance, but was identified in a VLSM of performance on two-proposition sentences compared to one-proposition sentences, suggesting that its role in sentence comprehension may be as a working memory or other task-related resource (Thothathiri et al. 2012).

Overall, large-scale lesion mapping studies have converged on the critical role of the posterior temporal-parietal region in sentence comprehension generally, but failed to substantially clarify the neural basis of agrammatic comprehension: Magnusdottir et al. implicated the ATL (for canonical > canonical) and Broca’s area (for non-canonical sentences alone) and Thothathiri et al. implicated only parietal-temporal regions for canonical, non-canonical, and canonical>non-canonical sentence measures. The discrepancy in findings between these two studies could be due to methodological differences. One difference is the testing of acute (Magnusdottir et al.) versus chronic stage (Thothathiri et al.) stroke patients. There are several possible differences in acute versus chronic stage testing that might affect VLSM results. For example, quantifiable areas of lesion are typically smaller in the acute stage than in the chronic stage (Birenbaum, Bancroft & Felsberg, 2011), leading to more focal brain regions being implicated in a particular diagnosis in acute patients (e.g. Ochfeld et al. 2010). Different brain regions also may be implicated for a given task in acute versus chronic stroke because of compensatory neural reorganization that has had more time to occur in chronic stroke (i.e. in chronic stroke different brain regions may support a task than in acute stroke) (Ochfeld et al. 2010; Kertesz 1997). A second difference is inclusion criteria (unselected vs. aphasia only): Magnusdottir et al. included left hemisphere stroke patients regardless of aphasia diagnosis; Thothathiri et al. only included left hemisphere stroke patients with a diagnosis of aphasia. The inclusion of only patients with aphasia could bias a sample toward larger lesion sizes and possibly exclude patients with more focal lesions and subclinical deficits, which in turn could lead to two different lesion patterns being implicated in a particular deficit. Including only patients with aphasia also potentially could bias results against Broca’s area in particular, given that restricted chronic damage to Broca’s area may present as non-aphasic dysarthria or apraxia of speech (Mohr, 1978; Alexander, Naeser & Palumbo 1990; Graff-Radford et al. 2014), and thus would not be represented in an aphasia only sample. Moreover, both studies used a difference score variable (canonical > non-canonical) that may yield misleading results with respect to agrammatic comprehension as it is classically defined: canonical performance near ceiling, non-canonical performance at chance level (Grodzinsky, 1990). Difference scores could pick up effects that are not as extreme as required to diagnose agrammatic comprehension (e.g., 10 percentage point differences) and would confound cases with correct performance at, for example, 90% canonical and 60% non-canonical (arguably a case of agrammatic comprehension) on the one hand and 60% canonical and 30% non-canonical on the other (clearly not such a case).

Another problem with previous large-scale studies of sentence comprehension generally and agrammatic comprehension more specifically is the use of sentence-picture matching tasks (e.g. Thothathiri et al. 2012; Magnusdottir et al. 2013). There is evidence from both large group and case studies that performance can dissociate across different types of sentence comprehension tasks, such as sentence-picture matching paradigms with whole sentence presentation, self-paced listening, plausibility judgments, and enactment (i.e. “object manipulation”) (Caplan, Michaud & Hufford, 2013, Caplan, DeDe & Michaud, 2006; Caplan, Waters & Hildebrandt, 1997; Gutman, DeDe, Michaud, Liu & Caplan, 2010; Cupples & Inglis, 1993; Tyler, Wright, Randall, Marslen-Wilson & Stamatakis, 2011). For example, Tyler et al.’s (2011) study of 14 aphasic patients compared areas of brain damage associated with performance on two types of sentence tasks, sentence picture matching and plausibility judgments. The sentence picture matching task identified several regions in which damage resulted in greater impairment on noncanonical compared to canonical sentences: Broca’s area, left posterior middle temporal gyrus, left superior temporal gyrus and the left supramarginal gyrus. The plausibility judgment task implicated a subset of these regions, namely Broca’s area and the left middle temporal gyrus (Tyler et al. 2011). Caplan, Michaud, Hufford & Makris (2016) have examined task-related differences in the regions of brain damage associated with impaired performance of aphasic patients in “off-line” (or after-the-fact assessment) versus “on-line” (or during sentence processing) tasks; off-line measures include sentence picture matching and object manipulation error rates; the on-line task measure is self-paced listening times. A broad summary of their findings is that damage to posterior superior temporal gyrus, inferior parietal regions such as supramarginal gyrus, angular gyrus, and Broca’s area contribute to agrammatic sentence comprehension to different degrees based on the task and type of noncanonical sentence used (passive, object-relative, reflexive, etc.).^1^ The implication of this kind of task-related variation is that task demands appear to be driving at least some of the results of mapping brain regions involved in sentence comprehension. Therefore, the use of only one task in previously published large-scale lesion studies are biased toward processes involved in that task.

The present study aimed to help resolve discrepancies regarding the contributions of Broca’s area and the ATL to sentence comprehension and specifically agrammatic comprehension. In addition to whole brain VLSM analyses and performance-based investigations of the lesions associated with agrammatic comprehension, we specifically target the relative contributions of Broca’s area and the ATL in sentence comprehension as these two regions are the primary foci for theories of the neural basis of sentence comprehension. Motivated by previous lesion work (e.g. Thothathiri et al. (2012) and Caplan et al. (2016)), we employ two different sentence comprehension tasks (sentence-picture matching and plausibility judgments) to help address task-related and cognitive resource contributions to sentence comprehension. There are very few lesion studies specifically addressing ATL function, particularly in relation to sentence processing. This may in part be due to the rarity of lesions from stroke in the anterior temporal lobe. Typically, ATL damage from stroke coincides with overall large left hemisphere damage and strokes restricted to the ATL are rare (Holland & Lambon Ralph, 2010). A more common patient group with focal ATL damage are people with temporal lobe epilepsy (TLE) who have undergone anterior temporal lobectomies. Patients after a lobectomy have been reported to have semantic, naming, and verbal fluency deficits (Ellis, Young & Critchley, 1989; Drane et al. 2009; Schwarz & Pauli, 2009; Frisk & Milner, 1990; Saykin et al. 1995; Janecek et al. 2013). Because these patients have medically intractable epilepsy, it is possible that their brains have abnormal functional organization prior to lobectomy including laterality of language networks (e.g. Swanson et al. 2002; Swanson, Sabsevitz, Hammeke & Binder, 2007; Hertz-Pannier et al. 2002) and any findings resulting from this population should be interpreted carefully. But, because of the rarity of natural ATL lesions (Holland & Lambon Ralph, 2010), the lobectomy population is worth investigating. Lexical deficits have been reported in this group (Glosser & Donofrio, 2001; Huang, Hayman-Abello, Hayman-Abello, Derry & McLachlan, 2014), but to our knowledge, the agrammatic pattern of sentence comprehension has not been yet been explicitly tested in lobectomy patients. Kho et al. (2008) report that 16 left lobectomy patients as a group (both pre-and post-surgery) did not perform significantly differently than controls on both active and passive sentences in a sentence-picture matching task. However, the aphasia literature indicates that the agrammatic performance pattern is more prominent for relative clause structures than passives (Berndt, Mitchum & Wayland, 1997) so perhaps more sensitive measures are needed to detect agrammatic comprehension patterns.

The present study took three approaches to address the relative contributions of Broca’s area, the ATL, and, in a more exploratory fashion, other areas to sentence processing. We first characterized the performance of patients with damage to Broca’s area (n=11) versus ATL damage (n=18) on two sentence comprehension tasks—sentence-to-picture matching and plausibility judgment—containing both canonical and noncanonical syntactic constructions. This allowed us to determine whether the agrammatic comprehension pattern of performance is causally related to damage in either of these regions. Second, across the entire sample, we identified patients with agrammatic comprehension as traditionally defined in the literature (Grodzinsky, 1989; Berndt, Mitchum & Haendiges, 1996), that is, performance at or below chance on the noncanonical sentences and significantly above chance on the canonical sentences; lesion distribution maps were then generated to identify areas of lesion overlap across patients. This was done separately for the two tasks to allow us to assess the typical lesion patterns associated with agrammatic comprehension and whether it varied as a function of task. We also investigated agrammatic comprehension by conducting a VLSM for each task to identify brain regions implicated in noncanonical sentence performance after variance due to canonical performance was regressed out. Finally, we conducted VLSMs for overall performance and for each sentence type in each task to identify the brain regions most implicated in comprehending the sentences more broadly.

If Broca’s area (or the ATL) is causally and differentially involved in processing noncanonical compared to canonical sentences, then:

1. patients with damage to Broca’s area (or the ATL) should perform at or near chance level on noncanonical items for both tasks and substantially better on canonical sentence structures (the agrammatic comprehension pattern).
2. patients with agrammatic comprehension on one or both tasks should have maximal lesion overlap in Broca’s area (or the ATL).
3. Noncanonical sentence comprehension, once variability due to canonical performance is controlled for, should implicate Broca’s area (or the ATL).
4. whole sample VLSM analyses should implicate Broca’s area (or the ATL) for noncanonical but not (or less so) for canonical sentences.

None of these expectations were borne out in the data. Instead, the data overall point to temporal/temporal-parietal areas as the primary region implicated in agrammatic comprehension and in sentence comprehension more generally.

## Methods

### Participants

141 patients were recruited via the Multi-site Aphasia Research Consortium (MARC) as part of an ongoing research program. Sixty-six of these patients were included in the present study based on the following criteria: (i) a chronic focal (6 months or more post-onset) lesion due to a stroke in the left hemisphere (n= 48; 17 female, mean age = 62 years, range = 31-84) or a unilateral left temporal lobectomy (n= 18; 10 female, mean age = 46, range = 29-62),^2^ (ii) no significant anatomical abnormalities other than the signature lesion of their vascular event (or evidence of surgery for the seizure subjects) nor signs of multiple strokes, and (iii) were administered both of the comprehension tasks of interest. Table 1 contains additional characteristics of the patients. All stroke patients had no prior history of psychological or neurological disease. The temporal lobectomy patients all had a seizure disorder that required lobectomy surgery for the treatment of their seizures. Patients were not selected for aphasia. Patients were all native speakers of English and the vast majority was strongly right-handed as determined by the Edinburgh Handedness Scale: two patients were left-handed and five were ambidextrous. Overall language abilities were assessed in each subject using the Western Aphasia Battery (WAB: Kertesz, 1982) and/or portions of the Boston Diagnostic Aphasia Examination (BDAE: Goodglass & Kaplan 1983) and clinician estimations. The aphasia subtypes present in the sample were as follows: 4 Wernicke’s, 7 Broca’s, 4 Conduction, 7 Anomic, 2 mild Broca’s / anomic, and 1 mixed nonfluent. The remainder of the patients (n=41, including all of the lobectomy patients) did not present with clinically significant language impairments on these measures. Informed consent was obtained from each patient prior to participation in the study, and all procedures were in compliance with the Code of Ethics of the World Medical Association and approved by the Institutional Review Boards of UC Irvine, San Diego State University, University of Southern California, University of Iowa and Arizona State University.

**Table 1.** Patient characteristics. “A” patients are in the ATL group, “B” patients are in the Broca’s area group, and “L” patients are additional left hemisphere patients included in the whole-brain analyses.

| ID | Gender | Age (Years) | Years post-stroke | Aphasia Subtype |
| --- | --- | --- | --- | --- |
| A01 | F | 42 | 2 | none |
| A02 | F | 43 | 4 | none |
| A03 | F | 61 | 6 | none |
| A04 | F | 59 | 6 | none |
| A05 | M | 53 | 5 | none |
| A06 | M | 44 | 2 | none |
| A07 | M | 29 | 3 | none |
| A08 | M | 47 | 14 | none |
| A09 | F | 39 | 5 | none |
| A10 | F | 61 | 1 | none |
| A11 | M | 49 | 4 | none |
| A12 | M | 31 | 3 | none |
| A13 | F | 34 | 3 | none |
| A14 | F | 54 | 1 | none |
| A15 | F | 47 | 10 | none |
| A16 | M | 62 | 0.5 | none |
| A17 | F | 39 | 6 | none |
| A18 | M | 35 | 2 | none |
| B01 | F | 61 | 13 | Broca's |
| B02 | M | 78 | 6 | Broca's |
| B03 | M | 59 | 4 | none |
| B04 | F | 57 | 10 | none |
| B05 | M | 63 | 4 | none |
| B06 | M | 68 | 1 | anomic |
| B07 | M | 61 | 13 | anomic |
| B08 | M | 72 | 1 | Broca's |
| B09 | M | 60 | 0.8 | none |
| B10 | M | 66 | 6 | anomic |
| B11 | F | 70 | 3 | conduction |
| L01 | F | 54 | 28 | none |
| L02 | M | 66 | 17 | none |
| L03 | F | 75 | 17 | none |
| L04 | F | 75 | 14 | none |
| L05 | M | 64 | 8 | Wernicke's |
| L06 | F | 67 | 9 | anomic |
| L07 | M | 43 | 4 | none |
| L08 | F | 79 | 31 | Wernicke's |
| L09 | F | 68 | 6 | none |
| L10 | M | 67 | 6 | none |
| L11 | M | 67 | 6 | none |
| L12 | F | 56 | 10 | none |
| L13 | M | 53 | 4 | anomic |
| L14 | M | 59 | 7 | none |
| L15 | M | 57 | 3 | none |
| L16 | M | 66 | 1 | conduction |
| L17 | M | 84 | 5 | anomic |
| L18 | M | 56 | 5 | none |
| L19 | M | 68 | 2 | conduction |
| L20 | M | 58 | 1 | Wernicke's |
| L21 | F | 64 | 2 | anomic |
| L22 | M | 68 | 5 | none |
| L23 | M | 60 | 0.5 | conduction |
| L24 | M | 72 | 3 | none |
| L25 | F | 34 | 2 | none |
| L26 | M | 57 | 3 | none |
| L27 | F | 55 | 3 | none |
| L28 | F | 81 | 2 | none |
| L29 | M | 51 | 2 | none |
| L30 | M | 48 | 9 | mixed nonfluent |
| L31 | F | 60 | 15 | Broca |
| L32 | M | 60 | 2 | Broca |
| L33 | M | 57 | 2 | mild Broca's /anomic |
| L34 | M | 47 | 6 | mild Broca's /anomic |
| L35 | F | 31 | 10 | Broca's |
| L36 | M | 67 | 1 | Wernicke's |
| L37 | M | 51 | 6 | Broca's |

### Materials

Patients completed two sentence comprehension tasks as part of the MARC test battery: the SOAP Test (a test of syntactic complexity; Love & Oster, 2002) and a plausibility judgment task. The two tasks are described below.

#### SOAP syntactic test

The SOAP is described in detail in Love and Oster (2002). Briefly, the SOAP is a sentence-picture matching task consisting of 40 trials, each of which involves a sentence being read to the subject, and the presentation of three colored drawings. The task is to point to the picture that corresponds to the sentence. There is a target (correct picture), a thematic role reversal foil and an unrelated foil. There are ten trials of each of the following sentence types: active, passive, subject-relative and object-relative. See Table 2 for examples. Proportion correct for each sentence type was calculated.

**Table 2.** Sentence-picture matching task sentence examples (for full list see Love & Oster, 2002).

|  |  |
| --- | --- |
| Active: | The girl in the green shirt chases the small boy. |
| Passive: | The boy in the green shirt is chased by the girl. |
| Subject-relative: | The boy that chases the girl is wearing a green shirt. |
| Object-relative: | The boy that the girl chases is wearing a green shirt. |

#### Plausibility judgments

The plausibility judgment task consists of 80 sentences (20 each of passive, active, subject-relative and object-relative; see Table 3). The sentences ranged in length from 8-10 words. The object-relative and subject-relative sentences were statistically indistinguishable regarding number of words (M = 10.7, sd =.47). Sentences were presented via headphones. 50% of sentences were plausible (could happen in the real world); 50% were implausible (see sentences in bold in Table 2). Patients were instructed to respond “yes” or “no” after listening to each sentence to indicate if the sentence was “plausible” that is, whether it “made sense”. Three practice trials containing active sentences were first presented to ensure that each patient understood the task.

**Table 3.**
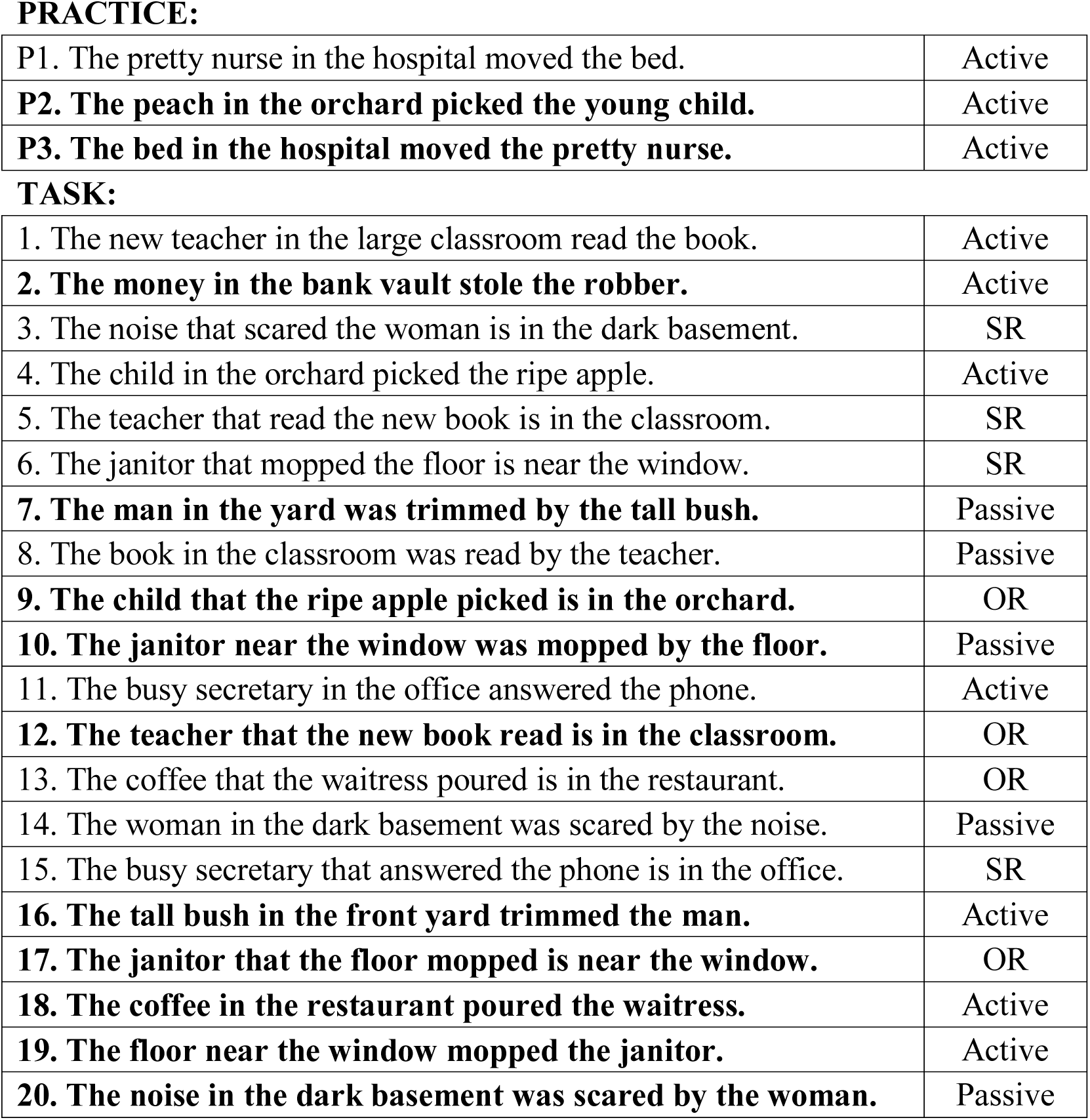
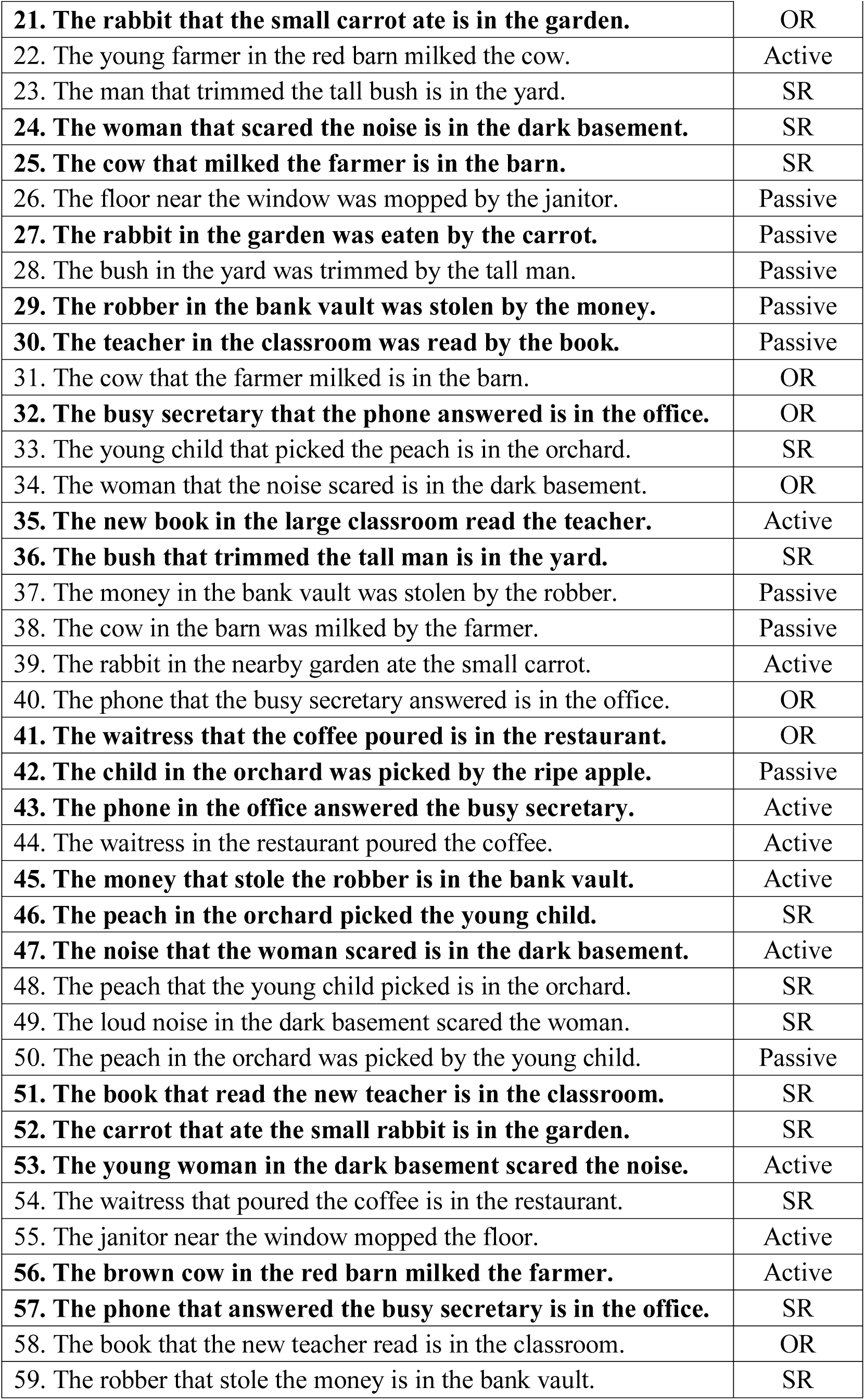
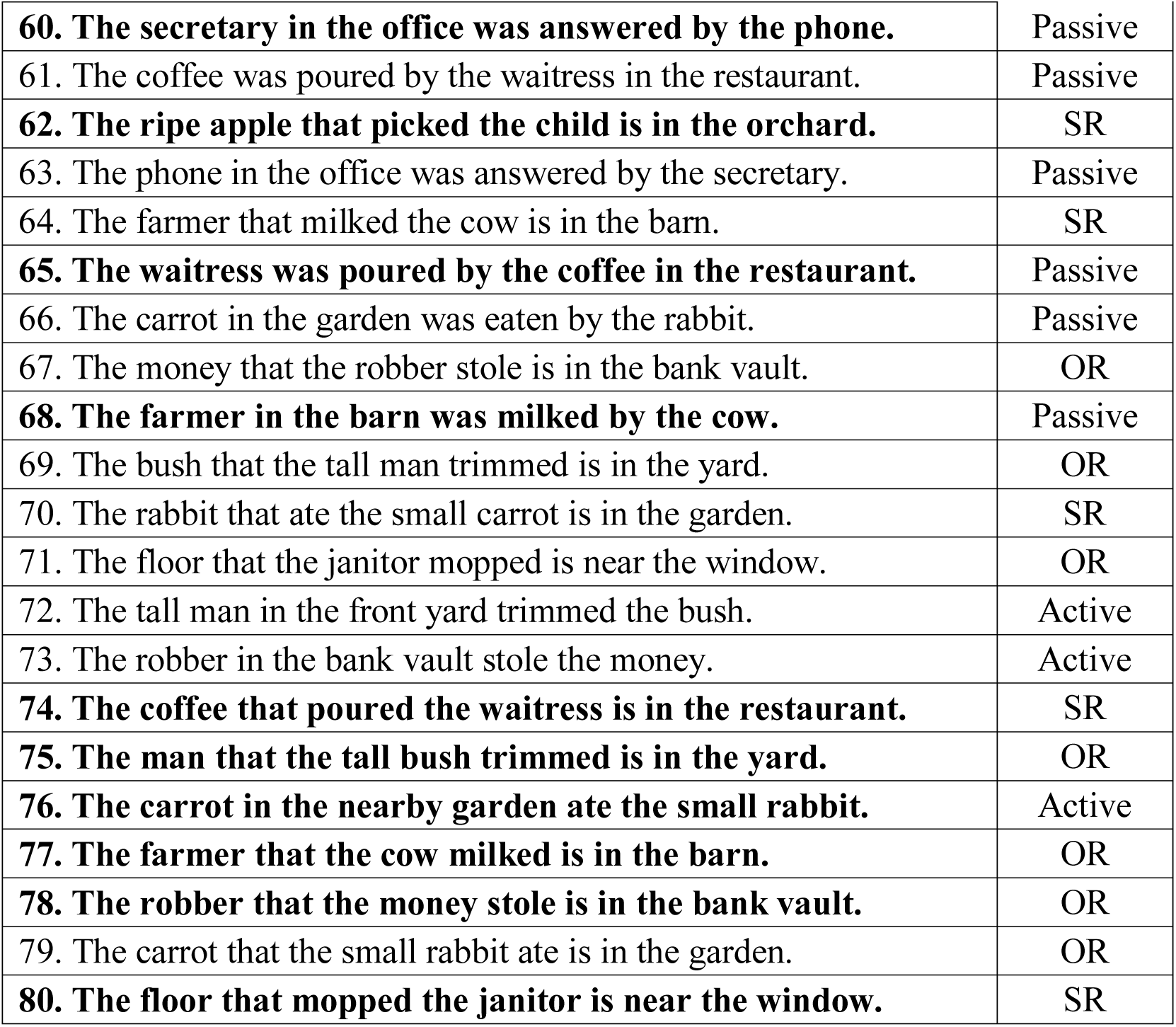
Plausibility judgment task sentences and sentence types. Implausible sentences are in bold.

**PRACTICE:**
|  |  |
| --- | --- |
| P1. The pretty nurse in the hospital moved the bed. | Active |
| **P2. The peach in the orchard picked the young child.** | Active |
| **P3. The bed in the hospital moved the pretty nurse.** | Active |
| **TASK:** |  |
| 1. The new teacher in the large classroom read the book. | Active |
| **2. The money in the bank vault stole the robber.** | Active |
| 3. The noise that scared the woman is in the dark basement. | SR |
| 4. The child in the orchard picked the ripe apple. | Active |
| 5. The teacher that read the new book is in the classroom. | SR |
| 6. The janitor that mopped the floor is near the window. | SR |
| **7. The man in the yard was trimmed by the tall bush.** | Passive |
| 8. The book in the classroom was read by the teacher. | Passive |
| **9. The child that the ripe apple picked is in the orchard.** | OR |
| **10. The janitor near the window was mopped by the floor.** | Passive |
| 11. The busy secretary in the office answered the phone. | Active |
| **12. The teacher that the new book read is in the classroom.** | OR |
| 13. The coffee that the waitress poured is in the restaurant. | OR |
| 14. The woman in the dark basement was scared by the noise. | Passive |
| 15. The busy secretary that answered the phone is in the office. | SR |
| **16. The tall bush in the front yard trimmed the man.** | Active |
| **17. The janitor that the floor mopped is near the window.** | OR |
| **18. The coffee in the restaurant poured the waitress.** | Active |
| **19. The floor near the window mopped the janitor.** | Active |
| **20. The noise in the dark basement was scared by the woman.** | Passive |
| **21. The rabbit that the small carrot ate is in the garden.** | OR |
| 22. The young farmer in the red barn milked the cow. | Active |
| 23. The man that trimmed the tall bush is in the yard. | SR |
| **24. The woman that scared the noise is in the dark basement.** | SR |
| **25. The cow that milked the farmer is in the barn.** | SR |
| 26. The floor near the window was mopped by the janitor. | Passive |
| **27. The rabbit in the garden was eaten by the carrot.** | Passive |
| 28. The bush in the yard was trimmed by the tall man. | Passive |
| **29. The robber in the bank vault was stolen by the money.** | Passive |
| **30. The teacher in the classroom was read by the book.** | Passive |
| 31. The cow that the farmer milked is in the barn. | OR |
| **32. The busy secretary that the phone answered is in the office.** | OR |
| 33. The young child that picked the peach is in the orchard. | SR |
| 34. The woman that the noise scared is in the dark basement. | OR |
| **35. The new book in the large classroom read the teacher.** | Active |
| **36. The bush that trimmed the tall man is in the yard.** | SR |
| 37. The money in the bank vault was stolen by the robber. | Passive |
| 38. The cow in the barn was milked by the farmer. | Passive |
| 39. The rabbit in the nearby garden ate the small carrot. | Active |
| 40. The phone that the busy secretary answered is in the office. | OR |
| **41. The waitress that the coffee poured is in the restaurant.** | OR |
| **42. The child in the orchard was picked by the ripe apple.** | Passive |
| **43. The phone in the office answered the busy secretary.** | Active |
| 44. The waitress in the restaurant poured the coffee. | Active |
| **45. The money that stole the robber is in the bank vault.** | Active |
| **46. The peach in the orchard picked the young child.** | SR |
| **47. The noise that the woman scared is in the dark basement.** | Active |
| 48. The peach that the young child picked is in the orchard. | SR |
| 49. The loud noise in the dark basement scared the woman. | SR |
| 50. The peach in the orchard was picked by the young child. | Passive |
| **51. The book that read the new teacher is in the classroom.** | SR |
| **52. The carrot that ate the small rabbit is in the garden.** | SR |
| **53. The young woman in the dark basement scared the noise.** | Active |
| 54. The waitress that poured the coffee is in the restaurant. | SR |
| 55. The janitor near the window mopped the floor. | Active |
| **56. The brown cow in the red barn milked the farmer.** | Active |
| **57. The phone that answered the busy secretary is in the office.** | SR |
| 58. The book that the new teacher read is in the classroom. | OR |
| 59. The robber that stole the money is in the bank vault. | SR |
| **60. The secretary in the office was answered by the phone.** | Passive |
| 61. The coffee was poured by the waitress in the restaurant. | Passive |
| **62. The ripe apple that picked the child is in the orchard.** | SR |
| 63. The phone in the office was answered by the secretary. | Passive |
| 64. The farmer that milked the cow is in the barn. | SR |
| **65. The waitress was poured by the coffee in the restaurant.** | Passive |
| 66. The carrot in the garden was eaten by the rabbit. | Passive |
| 67. The money that the robber stole is in the bank vault. | OR |
| **68. The farmer in the barn was milked by the cow.** | Passive |
| 69. The bush that the tall man trimmed is in the yard. | OR |
| 70. The rabbit that ate the small carrot is in the garden. | SR |
| 71. The floor that the janitor mopped is near the window. | OR |
| 72. The tall man in the front yard trimmed the bush. | Active |
| 73. The robber in the bank vault stole the money. | Active |
| **74. The coffee that poured the waitress is in the restaurant.** | SR |
| **75. The man that the tall bush trimmed is in the yard.** | OR |
| **76. The carrot in the nearby garden ate the small rabbit.** | Active |
| **77. The farmer that the cow milked is in the barn.** | OR |
| **78. The robber that the money stole is in the bank vault.** | OR |
| 79. The carrot that the small rabbit ate is in the garden. | OR |
| **80. The floor that mopped the janitor is near the window.** | SR |

Signal detection methods were used to determine how well subjects could discriminate plausible from implausible sentences. The signal detection measure *d’* was calculated for each sentence type. *D’* is a discriminability index indicating the distance between the means of the signal and the noise; d’ is reported in z-units, i.e. normalized standard deviation units (Swets, 1964). A larger *d’* indicates a larger separation between the number of “hits”, i.e. the number of times a plausible sentence was identified as plausible and the number of “false alarms,” i.e. the number of times an implausible sentence was identified as implausible. Thus, *d’* can be conceptualized as a performance measure that accounts for response biases. *D’* was used as the dependent measure for the plausibility judgments in the voxel-based lesion symptom analyses described below; however, proportion correct was used in the behavioral analyses to allow for comparison between the two sentence comprehension tasks.

### Imaging & lesion analyses

High-resolution MPRAGE or SPGR MRIs were acquired for all participants, except for four patients for whom a CT scan was acquired due to their incompatibilities with the MR environment (pacemaker, metal clip, etc.).

Lesion mapping was performed using MAP-3 lesion analysis methods (Damasio & Damasio, 2003) implemented on Brainvox software (Frank, Damasio & Grabowski, 1997). This method is described in detail elsewhere (Damasio, 2000). Briefly, the method entails the transfer of the lesioned brain areas as seen in the patient’s MRI/CT into the common space of a template brain. To do so, the template brain is resliced such that each slice maximally corresponds to each slice in the lesion’s native space, based on anatomical markers (e.g. sulci and subcortical structures). The lesion is then manually demarcated on the template brain’s corresponding slice respecting the identifiable landmarks. Each lesion map was completed by an individual with extensive training in this technique, and supervised by an expert neuroanatomist (HD). The template brain and resulting lesion maps were then transformed into Talairach space by Brainvox and resampled to a voxel size of 1mm^3^. The MAP-3 method has been shown to have high inter-and intra-rater reliability and in some cases higher accuracy than some automated methods (Fiez, Damasio & Grabowski, 2000; Pantazis, et al. 2010). Figure 1 depicts an overlay of all of the patients’ lesions maps on the template brain.

**Figure 1.**
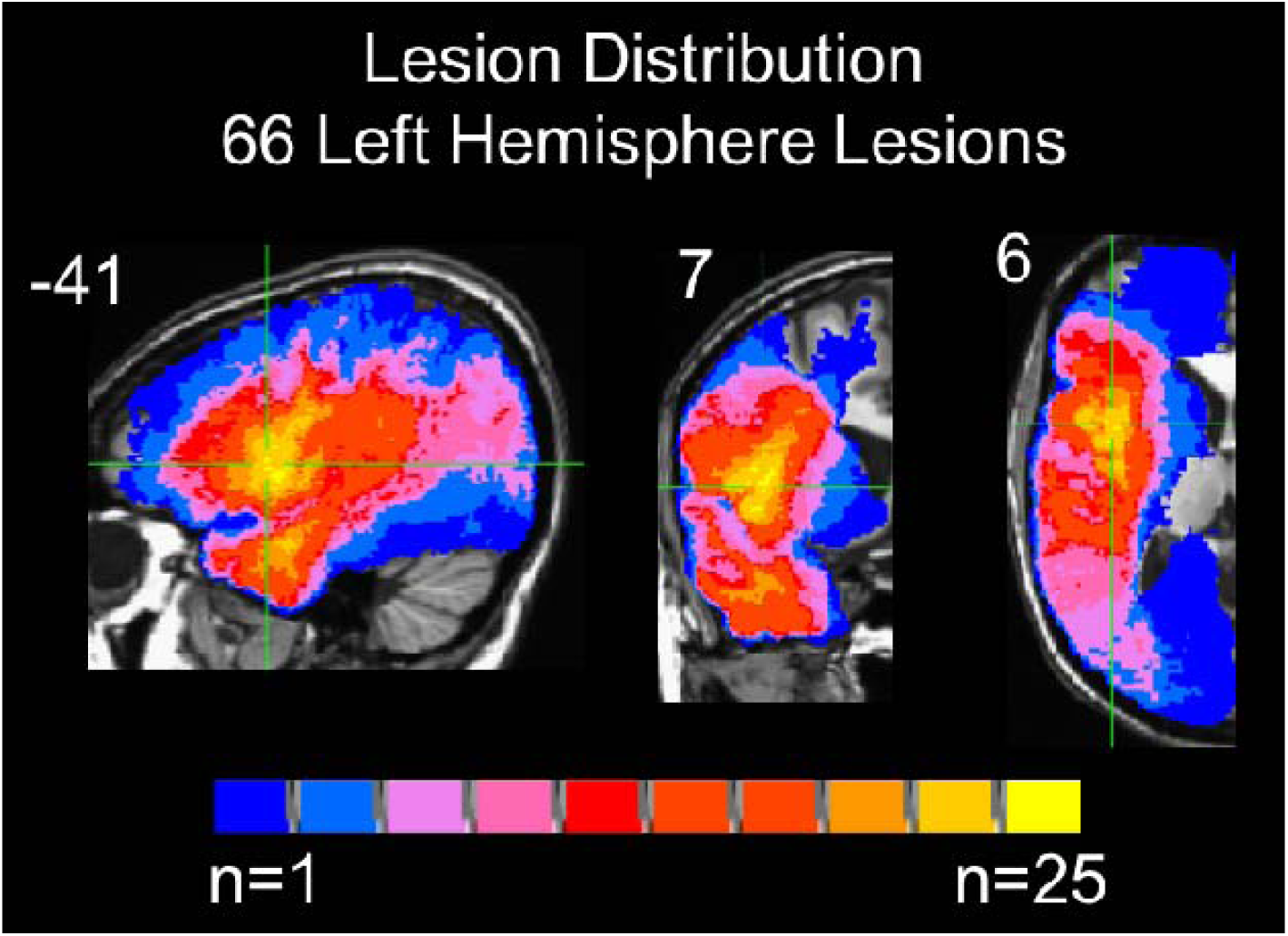
Overlap of all the patients’ lesions included in subsequent analyses (max overlap = 25). Coordinates are in Talairach space.

### Regions of Interest Analysis

Patients with damage anywhere in Broca’s area, defined as the posterior two-thirds of the left inferior frontal gyrus, i.e. the pars triangularis and pars opercularis (gray and/or white matter), without any temporal lobe involvement were identified for further study. Eleven patients met these criteria for inclusion in the Broca’s area damage group (Figure 2). The second group identified were patients with damage restricted to the left anterior temporal lobe, defined as any damage within any left temporal lobe gray and white matter anterior to the anterior edge of Heschl’s gyrus. 18 left temporal resections met these criteria (Figure 2). Sentence comprehension performance within and across these two groups was analyzed using a mixed-effect logistic regression for each task separately using PROC GLIMMIX in SAS software Version 9.4. Effects were estimated for group (Broca’s, ATL) and sentence type (OR, SR, passive, active) on response accuracy (correct vs. incorrect) in each task. Mixed-effect models were used as they provide a better fit of binomially distributed categorical data compared to a repeated measures analysis of variance and account for individual variability by computing a random intercept and slope per participant, allowing for a maximal model containing a maximum random effect structure (Jaeger, 2008; Barr, Levy, Scheepers & Tily, 2013). The fixed effects were group and sentence structure. Participants were treated as random effects. Pairwise comparisons were computed to investigate potential agrammatic sentence comprehension patterns within each group for each task. Multiple comparisons were corrected for by using Bonferroni-corrected p values, corrected across each group’s contrasts.

**Figure 2.**
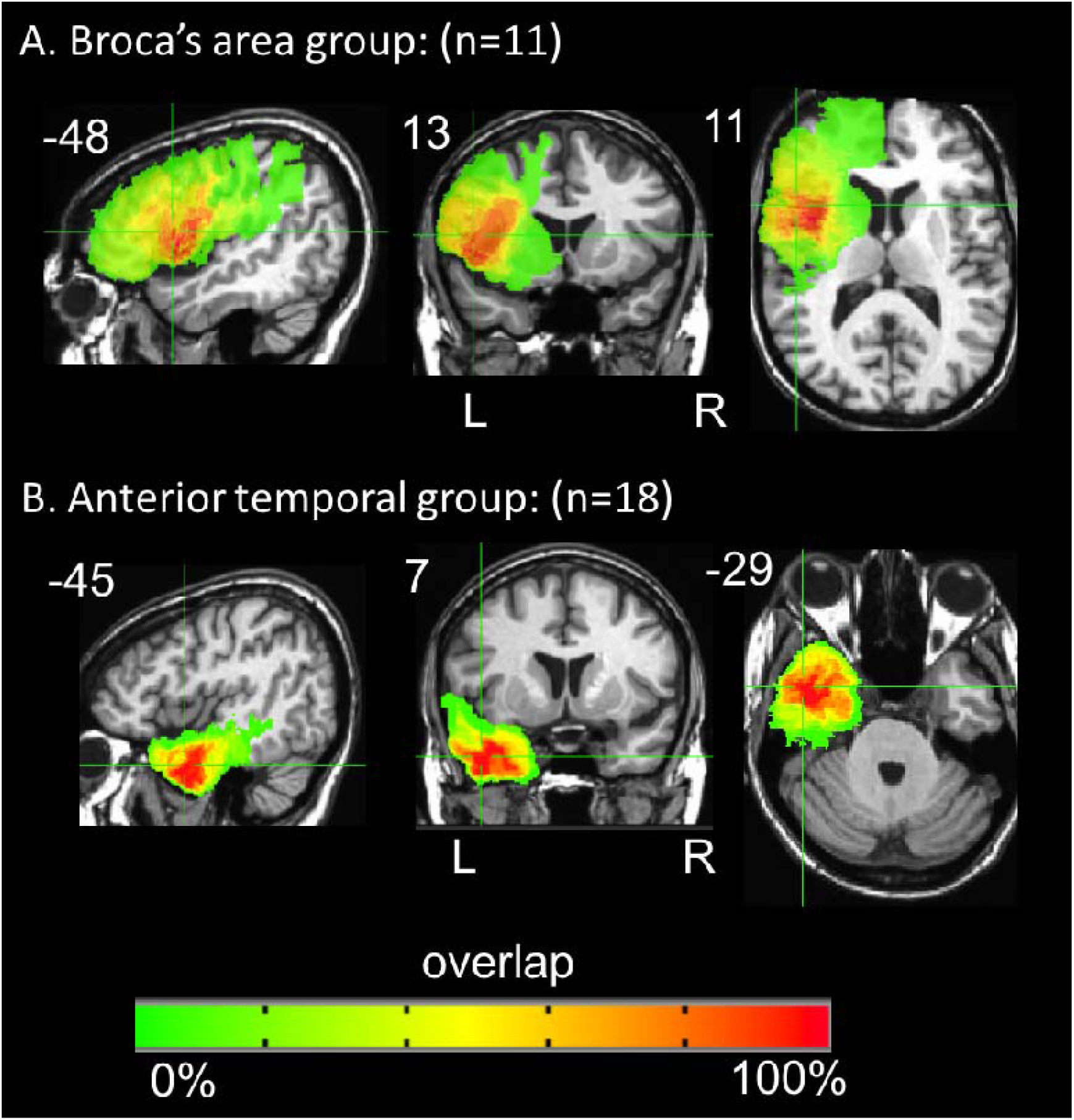
Overlap of the patients’ lesions in the (A) Broca’s area damage group (n=11, max overlap = 11) and (B) in the anterior temporal group (n=18, max overlap = 15). Coordinates are in Talairach space.

### Lesion Distributions of Agrammatic Comprehension

Agrammatic comprehension has traditionally been defined as performing significantly above chance on canonical sentences and at or below chance on noncanonical sentences (Grodzinsky, 1989; Berndt et al. 1996). It has been postulated that the strongest test of agrammatic comprehension is above chance performance on *all* canonical forms and at or below chance on *all* noncanonical forms (Grodzinsky, 1986). To examine the lesions associated with these criteria, binomial tests (p<.05, one-tailed) of performance in each task were used to identify patients in our sample who exhibited canonical sentence (i.e. subject-relative plus active) performance significantly above chance *and* noncanonical sentence (i.e. object-relative plus passive) performance not significantly above chance. Voxel-wise Liebermeister tests were computed using MRICron’s Non-Parametric Mapping (Rorden, Karnath & Bonilha, 2007) to identify regions of damage that are significantly associated with agrammatism. In addition, maps depicting the lesion distribution of the patients exhibiting agrammatic comprehension and of the patients not exhibiting agrammatic comprehension in each task were generated using AFNI’s 3dcalc command.

### Voxel-based lesion symptom mapping

In addition to the ROI-based and agrammatic comprehension lesion comparisons, voxel-based lesion symptom mapping (VLSM; Bates, et al. 2003) was used amongst all of the 66 left hemisphere patients (Figure 1) to identify voxels anywhere in the left hemisphere for which a t-test indicates that patients with damage in that voxel perform significantly different than patients who do not have damage in that voxel. For each task, a VLSM was calculated for performance on the noncanonical sentences (object-relative and passive sentences), with performance on the canonical (subject-relative and active) sentences as a covariate. In other words, variance due to canonical sentence performance was regressed out. These VLSMs should identify brain regions in which damage is associated with noncanonical < canonical performance, but they are not mathematically equivalent to VLSMs of canonical – noncanonical performance. As discussed in the introduction, difference score VLSMs might introduce confounds making interpretation of results difficult; thus we used a covariate approach instead.

Separate VLSMs also were completed for each sentence type in each task to further explore any differences between lesion patterns associated with noncanonical versus canonical sentence comprehension. A voxel-wise threshold of p <.001 was used. Multiple comparisons were controlled for using a cluster size-based permutation method: in each voxel, patients’ behavioral scores were randomly reassigned 1000 times. For each permutation, the general linear model was refit, a statistical threshold of p <.001 identified significant voxels, and the size of the largest cluster of significant voxels was recorded. Voxels in the actual data were identified as significant if they reached a voxel-wise p value <.001 and were in a voxel cluster larger than 95% of the largest significant clusters identified in the permutated datasets. (This cluster-based permutation method threshold procedure is the “p <.001, corrected” referred to in the subsequent results section). Variance due to lesion size (i.e. number of voxels marked as lesion on the template brain) was regressed out of all VLSM analyses via an Analysis of Covariance (ANCOVA). Only voxels in which a minimum of 10% of patients (n=7) had damage were included in the VLSM analyses.

### Response bias

The existence of task effects in sentence comprehension paradigms raises questions about the role of non-linguistic functions in performance. One potentially relevant measure, readily calculable from our plausibility task, is response bias. Frontal lobe regions have been previously implicated in decision-making processes in sentence comprehension tasks (Caplan, Alpert, Waters & Olivieri, 2008) and response bias in other speech tasks such as syllable discrimination (Venezia, Saberi, Chubb & Hickok, 2012; Baker, Blumstein & Goodglass, 1981; Hickok, Buchsbaum, Humphries & Muftuler, 2011). Thus, we examined the neural correlates of one measure of bias in the present study. Signal detection theory’s *C,* a representation of response bias, was calculated for each patient’s performance on the object-relative sentences in the plausibility judgment task (see Venezia et al. 2012 for a summary of signal detection theory as it relates to speech stimuli). The object-relative sentences were the focus of the response bias analysis because they elicited the largest range of performance and ceiling effects are unlikely. A VLSM was then generated to identify the areas of damage associated with greater response bias.

## Results

### Behavioral results – Broca’s area versus ATL damage

Performance of the Broca’s area damage group (n=11) and the left anterior temporal damage group (n=18) on both sentence tasks is presented in Figure 3 (solid bars). Within each task, accuracy for the two groups, i.e. patients with Broca’s area versus ATL damage, was carried out using mixed-model logistic regression with group (Broca’s vs. ATL) and sentence type (active, passive, SR, OR) as predictors.

**Figure 3.**
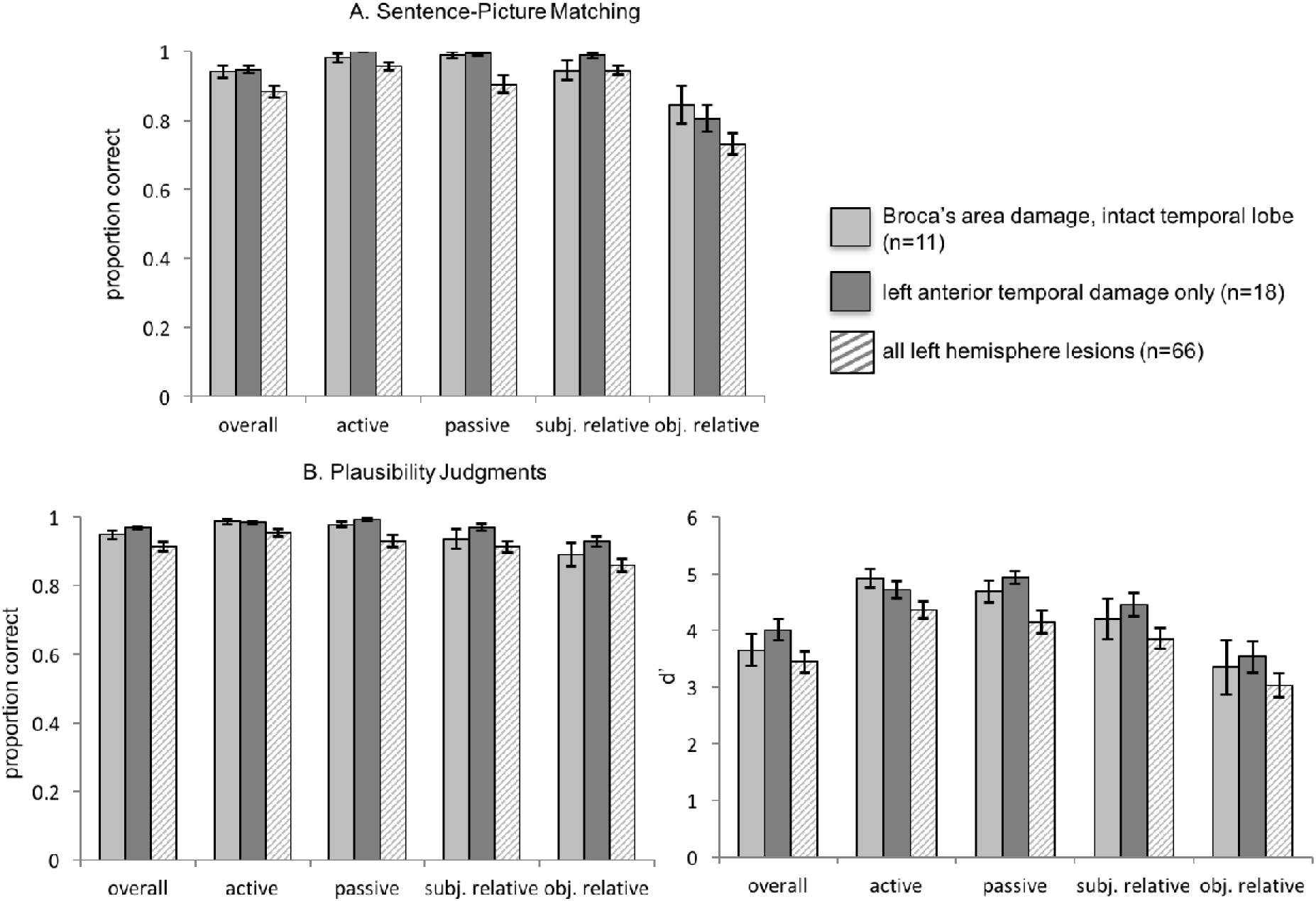
Graphs of mean group performance on the (A) sentence-picture matching task and on the (B) plausibility judgments. The light and dark gray solid bars represent the Broca’s and ATL groups, respectively. The striped bars represent performance of all LH subjects, e.g. all subjects included in the VLSM analyses. For the plausibility judgments, both proportion correct and d’ group mean performance are shown. Error bars represent standard error of the mean.

In the sentence-picture matching task, there was a significant main effect of sentence type (*F* = 12.5, *p*<.001); the main effect of group (*F* = 0, *p*=.99) and the group x sentence type interaction (*F*= 1.5, *p*=.23) were not significant. Note that the lack of a main effect of group indicates that the Broca’s and ATL damage groups do not exhibit significant differences in their sentence comprehension performance. Although the group x sentence type interaction was not significant, we conducted pairwise comparisons to further explore possible agrammatic comprehension patterns within each of the two groups because of our *a priori* prediction that if Broca’s area or the ATL is casually involved in agrammatic comprehension, one would expect an agrammatic comprehension pattern to be present in the corresponding group. Thus, we contrasted the parameter estimates for object-relative sentences versus subject-relative sentences, and for passive versus active sentences. In the Broca’s area group, there was no significant difference between object-relative and subject-relative performance (*β*= - 1.2, *SE* = 0.6, *p*=.22), and no significant difference between passive and active performance (*β*= 0.7, *SE* = 1.3, *p*=1). In the ATL group, object-relative performance was significantly lower than subject-relative performance (*β*= - 3.2, *SE*= 0.78, p=.0005), but there was no significant difference between passive and active performance (*β*= -14.6, *SE* = 1457.15, *p*=1.0).

The findings for the plausibility judgments are similar to the sentence-picture matching results: there was a significant main effect of sentence type (*F*= 12.4, *p*<.001); the main effect of group (*F*= 1.57, *p*=.22) and the group x sentence type interaction (*F*= 0.57, *p*=.63) were not significant. As in the sentence-picture matching task, we conducted pairwise comparisons based on our *a priori* predictions regarding agrammatic comprehension patterns within each group. The findings for the plausibility judgments were in line with those of the sentence-picture matching task: the Broca’s area group exhibited no significant difference between object-relative and subject-relative performance (*β*= -0.64, *SE*= 0.42, *p*=.77), as well as between passive and active performance (*β*= -0.53, *SE*= 0.77*, p*=1). In the ATL group, object-relative performance was lower than subject-relative performance, but this difference did not reach significance (*β*= -0.94, *SE*= 0.41*, p*=.14), and the difference between passive and active performance (*β*= 0.71, *SE*= 0.73*, p*=1.0) was not significant.

It is also noteworthy that the Broca’s area group and the ATL group both performed well above chance on the noncanonical sentences, i.e. the object –relative (SOAP mean correct = 85% for the Broca’s group and 81% for the ATL group; plausibility judgments mean correct = 89% for the Broca’s group and 93% for the ATL group) and passive sentences (SOAP mean correct = 99% for both the Broca’s and ATL groups; plausibility judgments mean correct = 98% for the Broca’s group and 99% for the ATL group). These results do not support the hypothesis that Broca’s area or the ATL is specifically linked to agrammatic comprehension.

Performance on each sentence type within each task for all 66 left hemisphere patients combined, i.e. all patients included in the VLSMs discussed below, are depicted in Figure 3 (striped bars). Performance of a group of 17 patients with right hemisphere damage due to stroke and no aphasia diagnoses is summarized in the supplemental material to provide a baseline of performance in each task.

### Results: Agrammatic Comprehension Lesion Distribution

Binomial tests to identify agrammatic comprehension indicate that in the sentence-picture matching task, 11 of the 66 left hemisphere patients performed significantly above chance on the canonical sentences and not significantly above chance on the noncanonical sentences (p<.05, one-tailed). In the plausibility judgment task, three left hemisphere patients performed significantly above chance on the canonical sentences and not significantly above chance on the noncanonical sentences (p<.05, one-tailed). At this point perhaps the most robust way to identify regions of damage associated with agrammatic comprehension as defined above would be to conduct voxel-wise Liebermeister tests (Rorden, Karnath & Bonilha, 2007) to determine if there are areas of damage that are significantly associated with the agrammatic group identified for each task. However, these tests yielded no significant results even at a liberal uncorrected threshold, likely in part due to insufficient power resulting from the small sample sizes of the agrammatic groups (n=11 and 3, respectively) (Rorden, Karnath & Bonilha, 2007). Thus we took a qualitative approach: maps of the lesion distributions for agrammatic and non-agrammatic comprehension in each task were generated to compare the lesions associated with agrammatic and non-agrammatic comprehension. These maps indicate the following: the patients with agrammatic comprehension in the sentence-picture matching have a maximum overlap (n=10 of 11) in the white matter underlying the left posterior superior and middle temporal gyri, with the center of mass being in the white matter underlying the left superior temporal gyrus, Talairach coordinates = -35 -48 17 (Figure 4A). Conversely, the patients who did not exhibit agrammatic comprehension in the sentence-picture matching task have maximum overlap in the ATL (n= 19 of 55, Talairach coordinates = -41 -6 -25); the second greatest area of overlap is in Broca’s area (n= 13 of 55) (Figure 4A). The three patients with agrammatic comprehension in the plausibility judgment task all have lesions overlapping in nearly the entire length of the left superior and middle temporal gyri and underlying white matter, extending into the inferior parietal lobe (Figure 4B). As in the sentence-picture matching task, patients who did not have agrammatic comprehension in the plausibility judgment task have maximum lesion overlap in the ATL (n= 22 of 63, Talairach coordinates = -43 0 -24) followed by an overlap of n= 17 of 63 in Broca’s area (Figure 4B).

**Figure 4.**
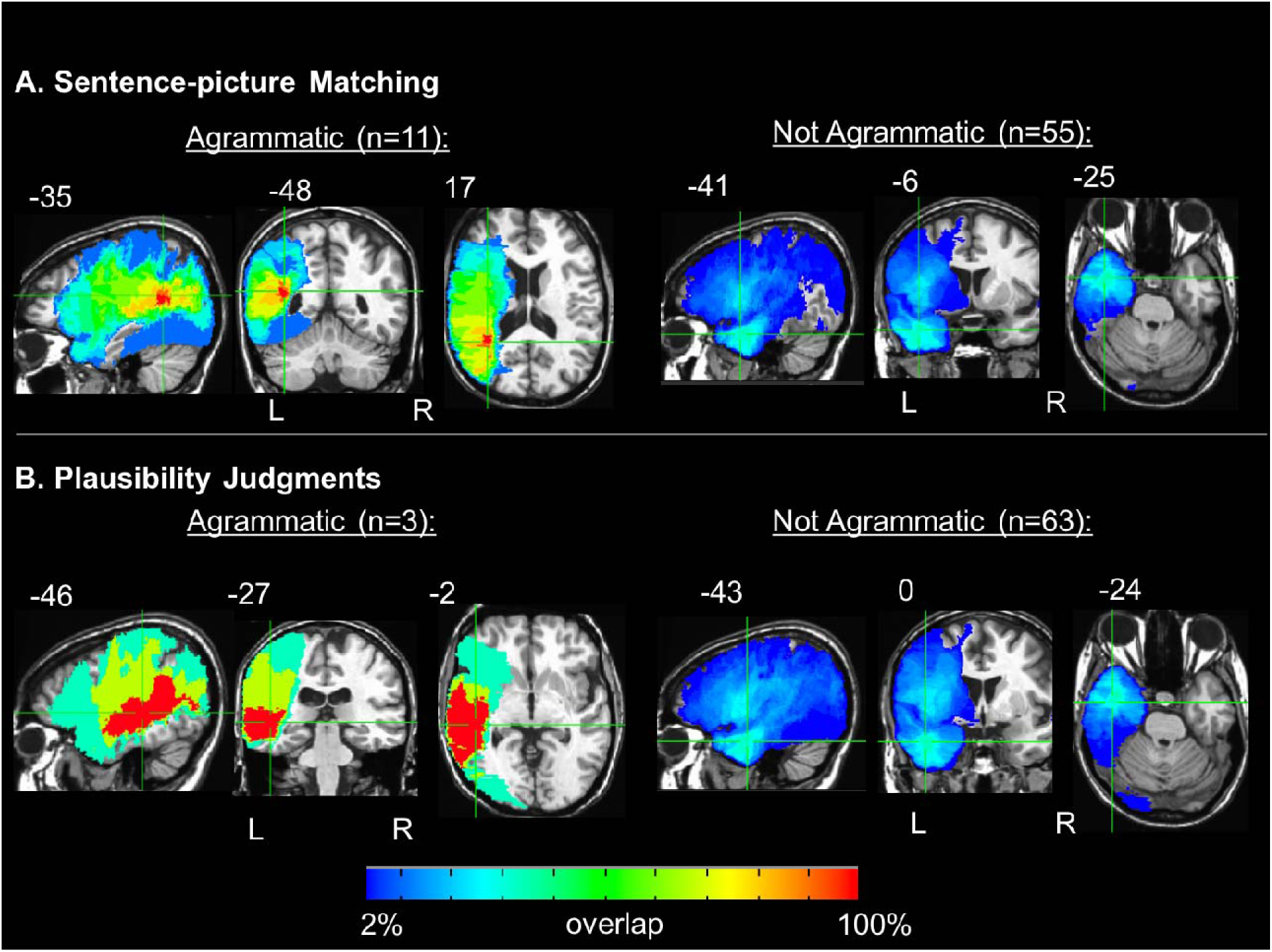
Orthogonal views of the lesion distributions for agrammatic and non-agrammatic performance on the (A) sentence-picture matching task and (B) plausibility judgments. Crosshairs are on regions of maximum overlap. Coordinates are in Talairach space.

The Broca’s area and ATL ROI analyses, and the lesion distributions of agrammatic comprehension, do not specifically implicate Broca’s area or the ATL in agrammatic comprehension. We now turn to whole brain analyses to examine the areas of damage associated with sentence comprehension deficits.

### VLSM Results: agrammatic comprehension

For both tasks, the VLSMs for performance on the noncanonical (OR and passive) sentences with canonical (SR and active) sentence performance as a covariate identified large clusters of voxels spanning the left superior temporal lobe and, to a lesser extent, the inferior parietal lobe: implicated regions include Heschl’s gyrus, the superior temporal gyrus, middle temporal gyrus, supramarginal gyrus, and underlying white matter (for sentence-picture matching peak *t* at Talairach coordinates -35 -52 14 in the white matter underlying the posterior superior temporal gyrus and number of voxels = 19650, for plausibility judgments peak *t* at Talairach coordinates - 54 - 26 5 and number of voxels = 13109, *p*<.001, corrected; Figure 5). The posterior boundary of the main clusters in each VLSM also extended into the middle occipital gyrus (>20 voxels).These findings align with the agrammatic comprehension lesion distribution results, implicating temporal and inferior parietal damage (but not frontal lobe damage) in agrammatic comprehension.

**Figure 5.**
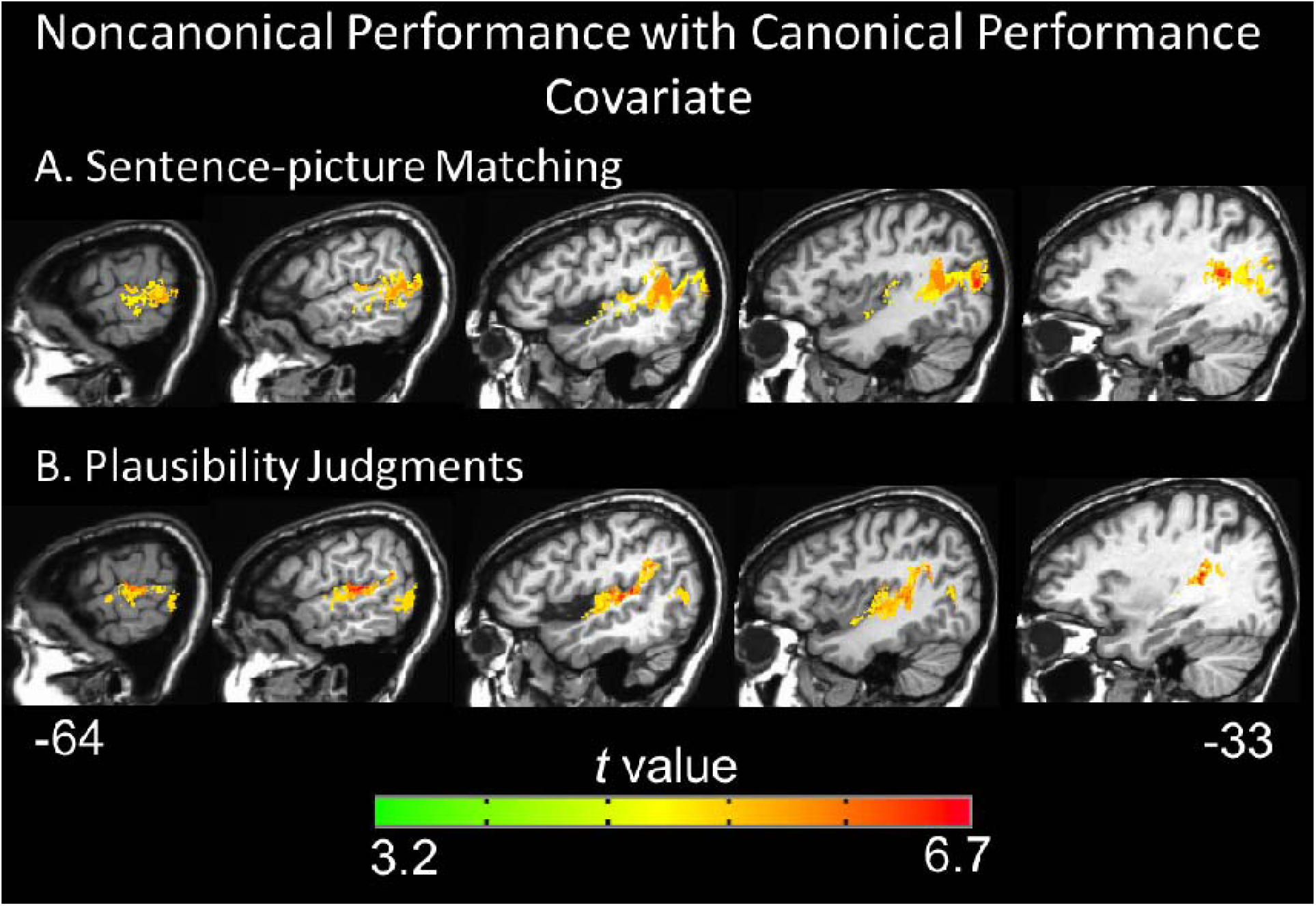
Representative sagittal slices depicting the VLSM results for performance on the noncanonical (object-relative and passive) sentences with canonical (subject-relative and active) sentence performance as a covariate regressed out on the (A) sentence-picture matching task and (B) plausibility judgments (voxel-wise height threshold = p<.001 and permutation-derived cluster threshold = p<.05). Coordinates are in Talairach space.

### VLSM Results: all sentence types

To further characterize the networks involved in sentence comprehension, we generated VLSMs for overall performance and performance within each sentence type, in each task, across all 66 left hemisphere patients. The VLSMs for overall performance on the sentence-picture matching task (proportion correct) and overall performance on the plausibility judgments (d’) both identified large clusters of voxels spanning the left superior temporal lobe and, to a lesser extent, the inferior parietal lobe: implicated regions include the superior temporal gyrus, middle temporal gyrus, angular gyrus, and underlying white matter (for sentence-picture matching peak *t* at Talairach coordinates -35 -46 13 and number of voxels = 63092, for plausibility judgments peak *t* at Talairach coordinates -45 -16 -5 and number of voxels = 40254 voxels, p<.001, corrected; Figure 6). The peak *t* value for the sentence-picture matching task was in the posterior superior temporal gyrus, while the peak t for the plausibility judgments was in superior temporal cortex just anterior to Heschl’s gyrus. Notably, the VLSMs for overall performance in both tasks identified significant voxels in the anterior temporal lobe (approximately in Brodmann areas 38 and anterior 22), but no frontal lobe voxels were significant. The VLSMs for each sentence type in each task all identify superior temporal and inferior parietal regions similar to the overall performance VLSMs; no frontal regions were significant for any sentence type in either task (Figures 7 and 8). These results indicate that a network of temporal and inferior parietal regions primarily support sentence processing.

**Figure 6.**
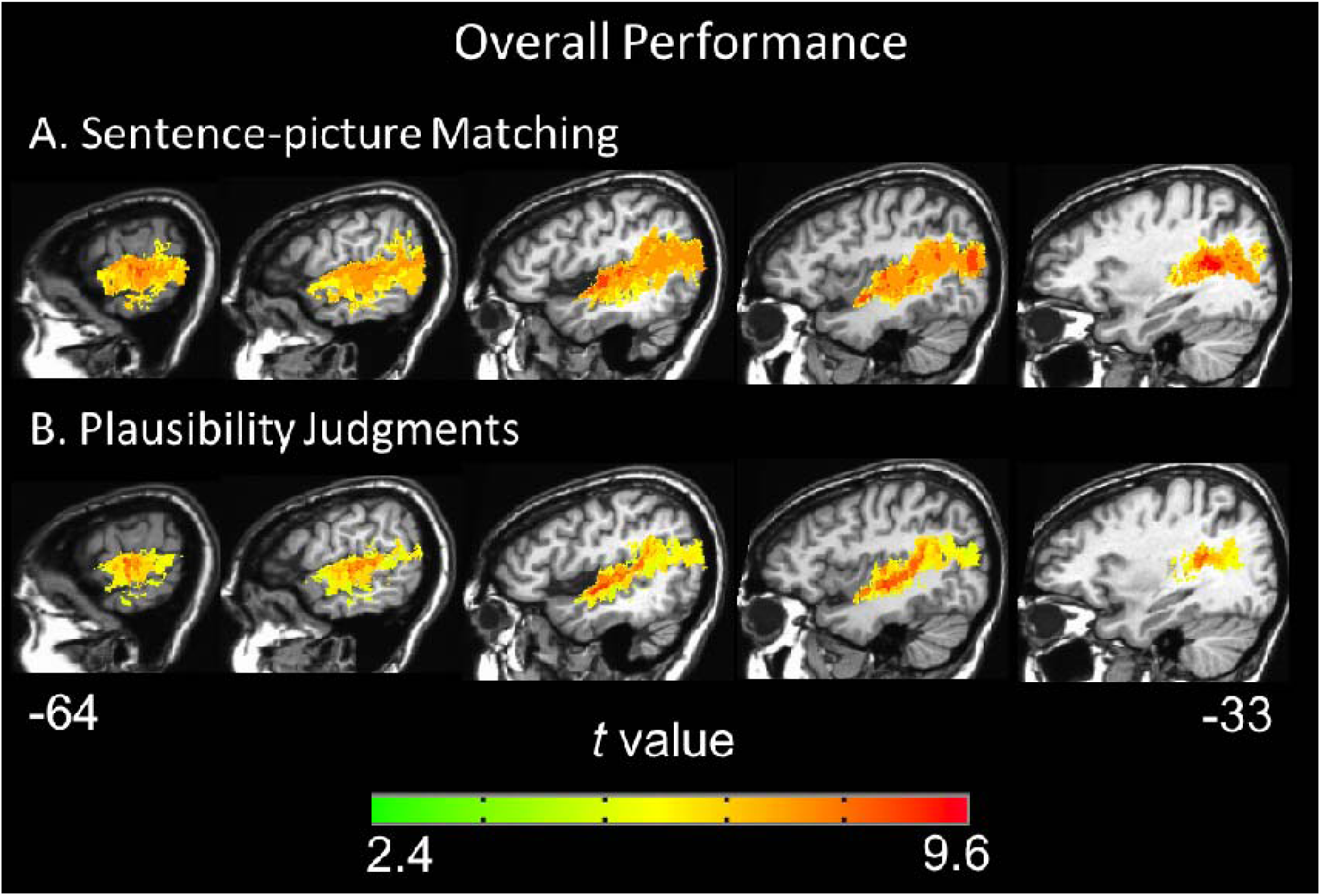
Representative sagittal slices depicting the VLSM results for overall performance on the (A) sentence-picture matching task and (B) plausibility judgments (voxel-wise height threshold = p<.001 and permutation-derived cluster threshold = p<.05). Coordinates are in Talairach space.

**Figure 7.**
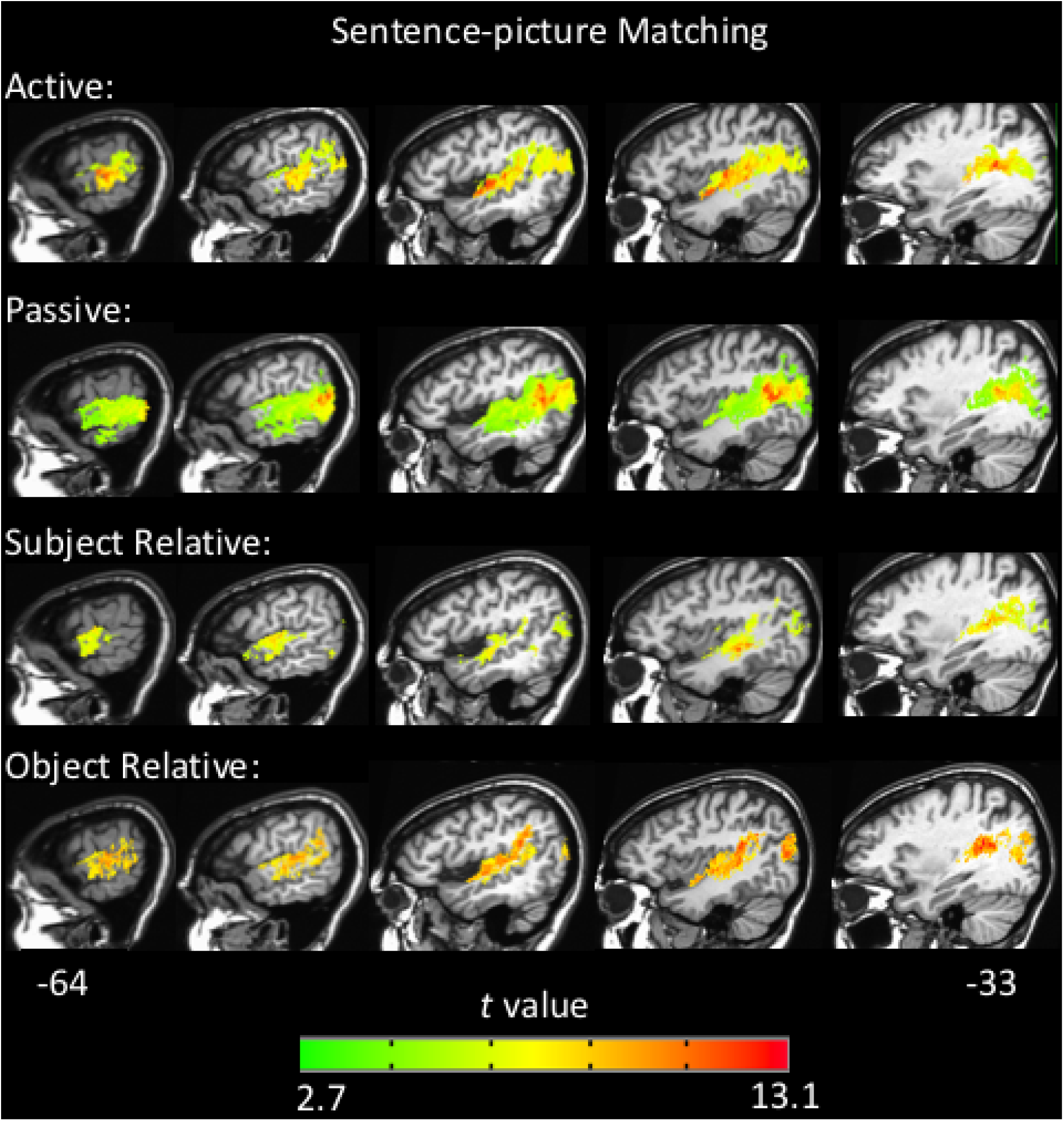
Representative sagittal slices depicting the VLSM results for performance within each sentence type in the sentence-picture matching task, (voxel-wise height threshold = p<.001 and permutation-derived cluster threshold = p<.05). Coordinates are in Talairach space.

**Figure 8.**
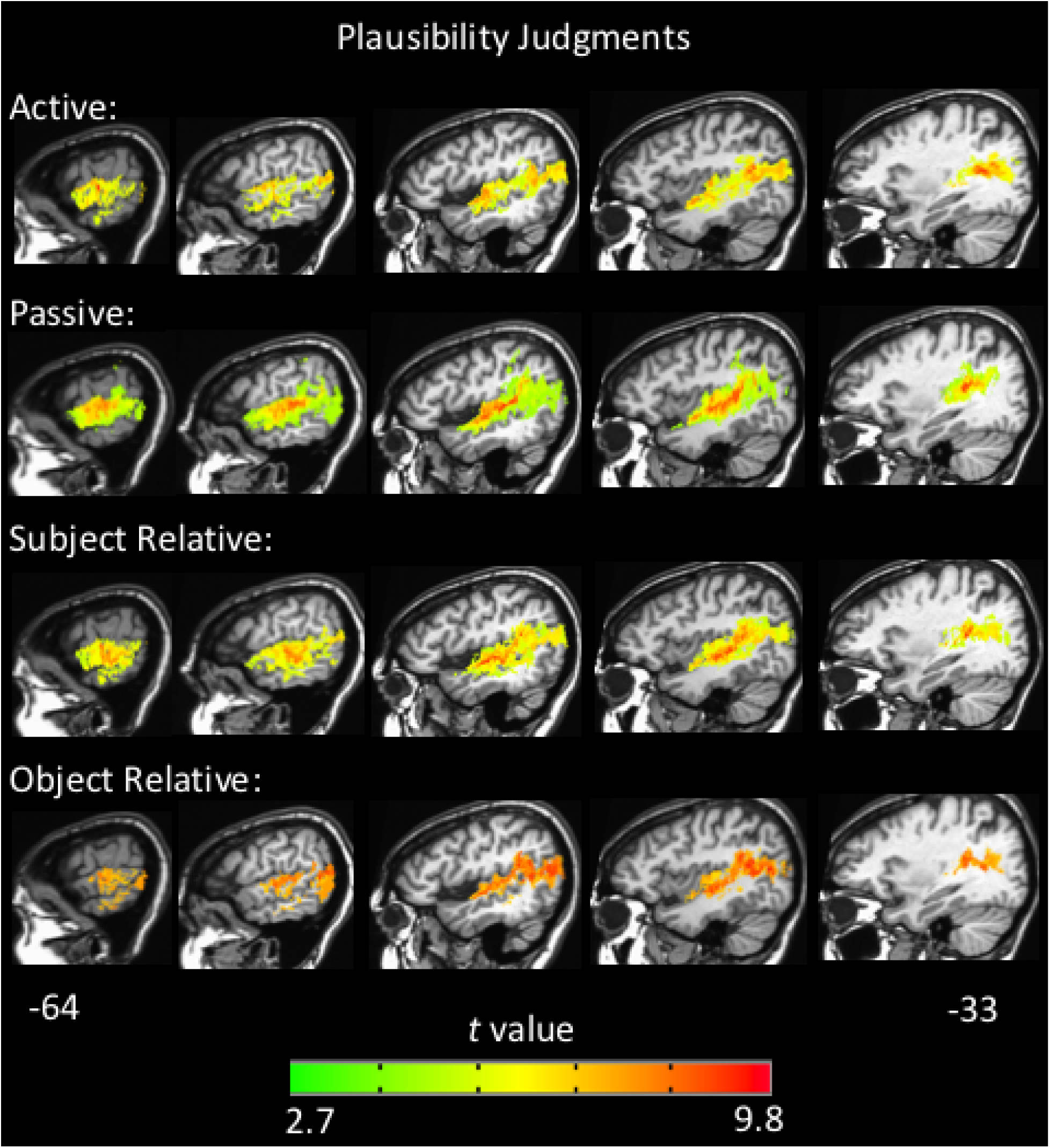
Representative sagittal slices depicting the VLSM results for performance within each sentence type in the plausibility judgment task, (voxel-wise height threshold = p<.001 and permutation-derived cluster threshold = p<.05). Coordinates are in Talairach space.

### Response bias & sentence comprehension

One representation of bias, *C*, was calculated for each patient’s performance on the OR sentences in the plausibility task. Of the 66 left hemisphere patients, 36 patients had a bias towards responding “yes” (i.e. yes, the sentence is plausible) while 8 were biased towards a “no” response. A VLSM was then generated to identify the areas of damage associated with greater response bias in either direction. The VLSM of bias identified a cluster predominately in the gray and white matter of the left inferior frontal gyrus (pars opercularis) and precentral gyrus, and extending into the insula and middle frontal gyrus (peak *t* at Talairach coordinates -50 10 5, number of voxels = 15264, p <.001, corrected, Figure 9). The absolute value of *C* was used as the variable of interest so that biases in both directions were included, but the results do not qualitatively differ by only examining the “yes” biases.

**Figure 9.**
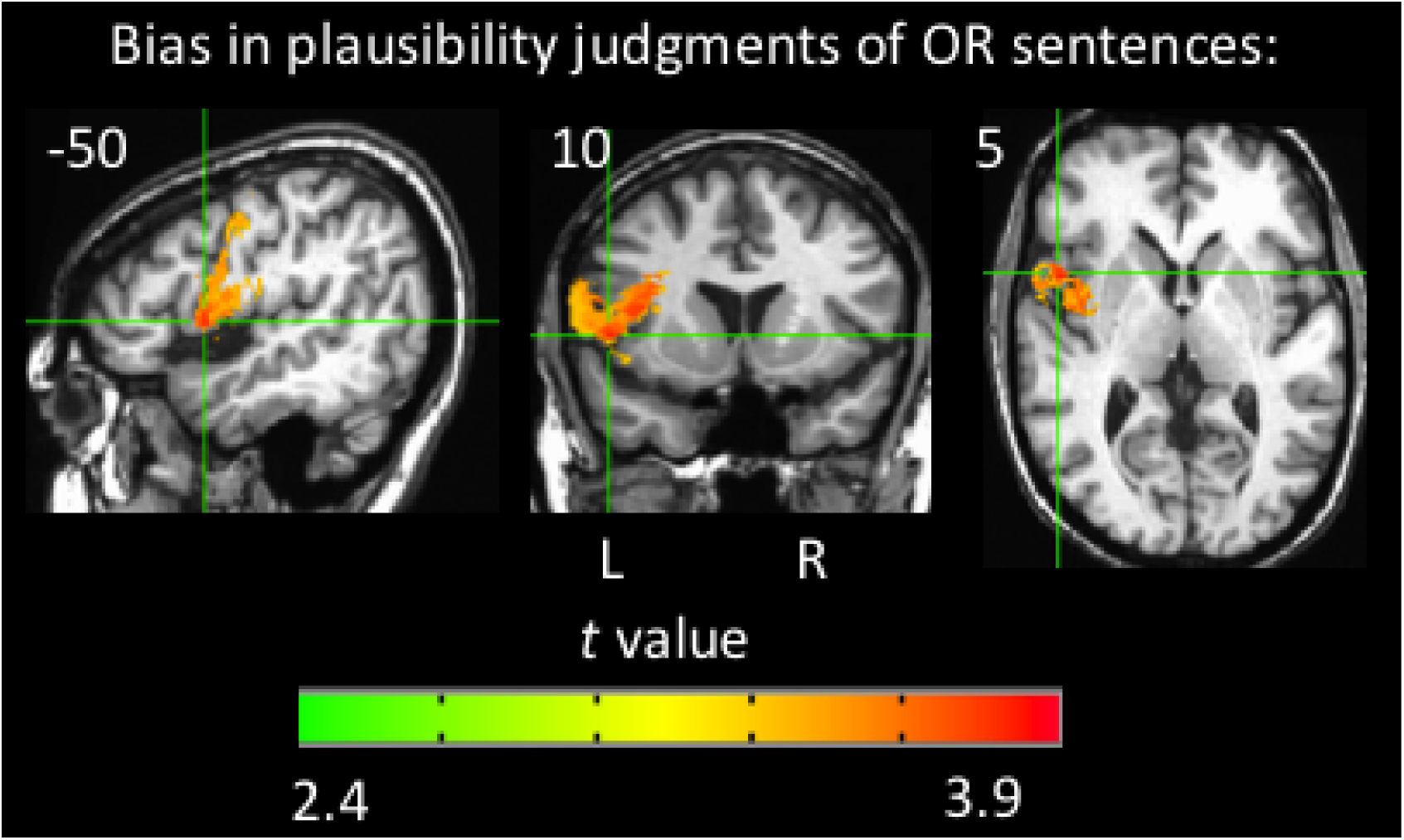
Orthogonal views of the VLSM results for bias in responses for OR sentences in the plausibility judgment task (voxel-wise height threshold = p<.001 and permutation-derived cluster threshold = p<.05). Crosshairs are on the peak *t* value. Coordinates are in Talairach space.

## Discussion

The present study investigated the neural substrates of agrammatic comprehension and sentence comprehension with a particular focus on the role of Broca’s area and the ATL. Patients with chronic, focal brain lesions completed two measures of sentence comprehension, a sentence picture matching task (the SOAP syntactic test; Love & Oster, 2002) and a plausibility judgment task. We implemented two different types of sentence tasks to be able to factor out any task-specific findings. Overall, our findings in lesion patients are highly consistent with previous large-scale studies of sentence deficits, which primarily implicate temporal and posterior parietal lobe networks (Magnusdottir et al. 2013; Dronkers et al. 2004; Thothathiri et al. 2012; Pillay et al. 2017). Our study adds to previous work in that we specifically assessed the contribution of Broca’s area and ATL damage.

### Sentence comprehension after damage to Broca’s area versus to the ATL

First, performance of patients with damage in Broca’s area (with the temporal lobe spared) were compared with performance in patients with damage to the ATL (with all other regions spared). Analyses comparing the Broca’s area and ATL groups revealed highly similar behavioral performance; there were no significant differences in performance between the two groups for any sentence type in either task. The Broca’s area group did not exhibit significantly lower performance for the noncanonical sentences than for the canonical sentences (contrasts of OR versus SR and passive versus active performances were not significant in either task). The ATL group did demonstrate significantly lower performance for OR versus SR sentences in the sentence-picture matching task, but in the plausibility judgment task this effect only approached significance (*p* =.14); the passive versus active contrast was not significant in either task for the ATL group. Both groups performed well above chance on the object-relative sentences on both tasks (> 80% correct on the SOAP task and d’ > 3 on the plausibility judgment task) in contrast to previous claims that damage to Broca’s area is responsible for agrammatic comprehension, i.e., near chance performance on object relatives (Grodzinsky, 1989, 2000). The finding that damage to Broca’s area is not reliably associated with agrammatic comprehension coincides with previous lesion work (Thothathiri et al. 2012; Magnusdottir et al. 2013; Caramazza, Capasso, Capitani & Miceli, 2005). The finding that the ATL is not strongly associated with agrammatic comprehension is at odds with the agrammatic comprehension findings that were localized to the ATL in Magnusdottir et al.’s study of acute stroke patients. However, neural reorganization needs to be considered when interpreting our findings in the ATL; our ATL group consists of temporal lobectomy patients who have likely experienced functional neural reorganization due to their medically intractable epilepsy and perhaps additionally after their temporal resection surgery.

### Lesion overlap in cases of agrammatic comprehension

Agrammatic comprehension (significantly above chance on canonical sentences, not significantly above chance on noncanonical sentences) in each task was predominantly associated with superior temporal and inferior parietal damage and not Broca’s area (or ATL) as standard models would predict. More specifically, the area of maximum overlap among patients with agrammatic comprehension was in the posterior STG. ATL damage was present in 2 of the 3 patients with agrammatic comprehension on the plausibility judgments and in 7 of 11 patients with agrammatic comprehension on the sentence-picturing matching task. Broca’s area damage was present in 1 of 3 patients with agrammatic comprehension on the plausibility judgments and in 4 of 11 patients with agrammatic comprehension on the sentence-picture matching task. However, the areas of greatest overlap in the non-agrammatic comprehension group in each task were the ATL (35% for each task) followed by Broca’s area (27% for plausibility judgments and 24% for sentence-picture matching). Thus, the agrammatic comprehension overlap maps provide evidence for loose relations at best between agrammatic comprehension, and ATL or Broca’s area damage. Based on lesion overlap analyses, then, we conclude that the posterior STG is a more critical site in producing agrammatic comprehension than either Broca’s area or the ATL.

### Voxel-Based Lesion Symptom Mapping of Non-Canonical Sentence Comprehension

The VLSMs investigating brain regions implicated in noncanonical sentence comprehension with canonical sentence comprehension performance as a covariate also identified superior temporal and inferior parietal regions in both tasks (Figure 5) consistent with the agrammatic comprehension lesion overlap findings (Figure 4). The implicated regions in the STG extended into areas anterior to Heschl’s gyrus: for the sentence-picture matching task 212 voxels were significant on the superior bank of the anterior STG, and for the plausibility judgment task 156 voxels were significant on the superior bank and lateral aspects of the anterior STG. The coordinates of the peak *t* values in the ATL for both of these VLSMs fall within the area of maximum overlap in the ATL in the agrammatic comprehension sentence-picture matching overlap map. The ATL regions identified by these VLSMs do overlap with the ATL regions found in some fMRI studies to be more activated in response to sentences than non-sentence stimuli (e.g. Friederici et al. 2010; Rogalsky et al. 2011). However, other fMRI findings implicate the middle temporal gyrus as well as the STG, and the activations extend more towards the temporal pole, extending into approximately Brodmann area 38, whereas our VLSM findings are more posterior and solely in approximately Brodmann area 22 (e.g. Humphries et al. 2006; Stowe et al. 1998; other contrasts in Rogalsky et al. 2011). We found no evidence of Broca’s area involvement in non-canonical sentence comprehension on either of the tasks.

To summarize, our VLSMs of non-canonical sentence comprehension primarily implicate posterior temporal and inferior parietal regions with only partial extension into ATL regions previously implicated in sentence processing and no evidence of Broca’s area involvement. This finding of temporal and parietal regions implicated in agrammatic comprehension is highly consistent with previous VLSM studies of agrammatic comprehension (Thothathiri et al. 2012; Magnusdottir et al. 2013; Pillay et al. 2017) as well as previous functional neuroimaging studies of sentence comprehension generally (Vandenberghe et al. 2002; Humphries et al. 2005, 2006, 2007; Snijders et al. 2009; Friederici, 2011). It also should be noted that the null results in Broca’s area in the present study are not a product of reduced sensitivity due to multiple comparison correction; the uncorrected results also do not implicate Broca’s area. Insufficient power also can be a concern when interpreting null findings. While possible, it is unlikely here given that the greatest lesion coverage of our sample is centered in Broca’s area (overlap of n =25 in the pars opercularis, plus an additional five subjects not contributing to this maximum overlap but who do have damage elsewhere in Broca’s area and/or its underlying white matter).

### Voxel-Based Lesion Symptom Mapping of Sentence Comprehension Generally

Whole brain VLSM analyses examining the relation between comprehension on the various sentence types for the two tasks in the entire left-hemisphere damaged sample showed similar patterns of lesion-symptom mappings. Specifically, all sentence types on both tasks implicated superior and middle temporal lobe regions including the superior temporal gyrus, middle temporal gyrus, and extended into inferior parietal cortex. These VLSM analyses of each sentence type all identified more anterior extension in the temporal lobe than that found in the non-canonical sentence analysis, which provides some support for the claim that the ATL is involved in basic combinatorial sentence processing (Humphries, Binder, Medler & Liebenthal, 2006; Rogalsky & Hickok, 2009; Rogalsky, Rong, Saberi & Hickok, 2011; Brennan & Pylkkanen, 2012; Brennan et al. 2012). The ATL regions identified here overlap with ATL regions previously implicated in sentence processing by fMRI studies (e.g. Friederici et al. 2010; Stowe et al. 1998; Rogalsky et al. 2011), although the lesion maps do not extend as anteriorly as some reported fMRI peak activations (e.g. Vandenberghe et al. 2002; Rogalsky & Hickok, 2009).

In contrast to the involvement of at least some portions of the ATL in sentence comprehension, Broca’s area was not implicated in these analyses, even for the sentence types (object relatives) that have been classically associated with damage to or activation of the IFG. What this finding suggests is that basic receptive processes in auditory language comprehension are supported by the temporal and posterior parietal lobes rather than frontal lobe regions (Bates et al. 2003).

### Interpretation, limitations, and comparison to previous work

Consistent with three recent large-scale studies from three different groups (Thothathiri et al. 2012; Magnusdottir et al. 2013; Pillay et al. 2017), the present study found no evidence for the involvement of Broca’s area in agrammatic comprehension in any of our analyses designed specifically to test the hypothesized linkage. Furthermore, no evidence was found for the role of Broca’s area in sentence processing more generally. The ATL was found to be more related to sentence processing generally than was Broca’s area, although the ATL’s link to agrammatic comprehension was weak in our sample (c.f. Magnusdottir, et al. 2013). The most consistently implicated region in agrammatic comprehension and sentence processing generally was the posterior superior temporal and inferior parietal regions, which have emerged recently as the core of the sentence processing network.

The finding that the superior temporal and inferior parietal region is more implicated in sentence comprehension impairments than Broca’s area present a paradox given the long-standing association between Broca’s aphasia and agrammatic comprehension (Caramazza & Zurif, 1976; Grodzinsky 1986, 2000). The paradox can be resolved, however, by noting two facts. The first is that agrammatic comprehension is not unique to Broca’s aphasia but has also been documented in conduction aphasia (Caramazza & Zurif, 1976), which is associated with posterior temporal-parietal lesions (Damasio & Damasio,1980; Buchsbaum, Baldo, Okada, Berman, Dronkers, D’Esposito & Hickok, 2012). The second fact is that Broca’s aphasia is not tied to Broca’s area alone (Mohr et al. 1978; Mohr, 1976; Fridriksson, Bonilha & Rorden, 2007) and more recently has been shown to involve damage to the pars opercularis portion of Broca’s area in combination with the posterior superior temporal lobe (Fridriksson, Fillmore, Guo & Rorden, 2015). Thus, classical lesion-symptom evidence is quite consistent with a posterior temporal and parietal focus for sentence processing including agrammatic comprehension.

Another paradox arises when considering the present results alongside of the functional imaging literature, which has frequently implicated Broca’s area in sentence processing (e.g. Bedny, Pascual-Leone, Dodell-Feder, Fedorenko & Saxe, 2011; Rogalsky et al. 2015; Blank, Balewski, Mahowald & Fedorenko, 2016). As mentioned in the introduction, a possible resolution of this paradox comes from hypotheses that Broca’s area supports performance on sentence comprehension tasks through more general cognitive functions such as working memory (Just, Carpenter, Keller, Eddy & Thulbom, 1996; Kaan & Swaab, 2002; Rogalsky, Matchin & Hickok, 2008; Pettigrew & Hillis, 2014) and/or cognitive control (Novick, Trueswell & Thompson-Schill, 2005). The present finding of a link between response bias and Broca’s area lesions may reflect some aspect of cognitive control or working memory impairment (Venezia et al. 2012) as both of these processes have in part been localized to Broca’s area (Novick et al. 2005; Smith & Jonides, 1997; Hickok et al. 2003). Neuroimaging studies of response selection (in which deficits can result in response bias) consistently implicate Broca’s area and adjacent prefrontal cortex (e.g. Goghari & MacDonald, 2009; Harding, Yucel, Harrison, Pantelis & Breakspar, 2015). In addition, working memory impairments due to inferior frontal damage may prevent the use of articulatory rehearsal to assist in the maintenance of a sentence that is difficult to comprehend, thereby causing the patient to choose the most likely response based on previous experience (Venezia et al. 2012) which in the case of sentences heard in everyday life, the sentence is plausible). These possibilities would explain Broca’s area activation in many sentence comprehension studies yet also explain why Broca’s area doesn’t always activate during sentence comprehension (e.g. Sptisyna et al. 2006; Humphries et al. 2005, 2006; Rogalsky et al. 2011): depending on the particular task demands, support functions mediated by Broca’s area will be recruited or not. This finding also coincides with Thothathiri et al.’s finding that damage to Broca’s area (specifically Brodmann area 44), and not temporo-parietal regions, is associated with decreased comprehension of two-proposition sentences compared to one-proposition sentences. Thothathiri et al. suggest that increased working memory demands of the two-proposition sentences compared to the one-proposition sentences explain this effect. Notably, several of the recent high-profile functional imaging studies implicating Broca’s area in sentence processing have included a memory task (Blank, Balewski, Mahowald & Fedorenko, 2016; Fedorenko, Hsieh, Nieto-Castanon, Whitfield-Gabrieli & Kanwisher, 2010; Fedorenko et al. 2011, Fedorenko, Duncan & Kanwisher, 2012a; Fedorenko, Nieto-Castanon & Kanwisher, 2012b, 2012c). The inclusion of a memory task likely engages working memory processes, including articulatory rehearsal, which robustly activates Broca’s area (Hickok, Buchsbaum, Humphries & Muftuler, 2003; Buchsbaum, Ye & D’Espositio, 2011; Buchsbaum & D’Esposito, 2008; Smith, Jonides, Marshuetz & Koeppe, 1998).

Another relevant factor that is often overlooked in interpreting functional activation studies of sentence processing is whether the stimuli are presented visually (written sentences) or auditorily. We have noticed that a disproportionate number of sentence processing studies that implicate Broca’s area have used written stimuli (Table 4). Given that reading involves articulatory processes to a greater extent than listening (Daneman & Newson, 1992; Slowiaczek & Clifton, 1980; Baddeley, Thomson & Buchanan, 1975), the use of written sentences in many past studies appears to have biased the functional imaging literature toward a Broca’s-area-centric perspective.

**Table 4.**
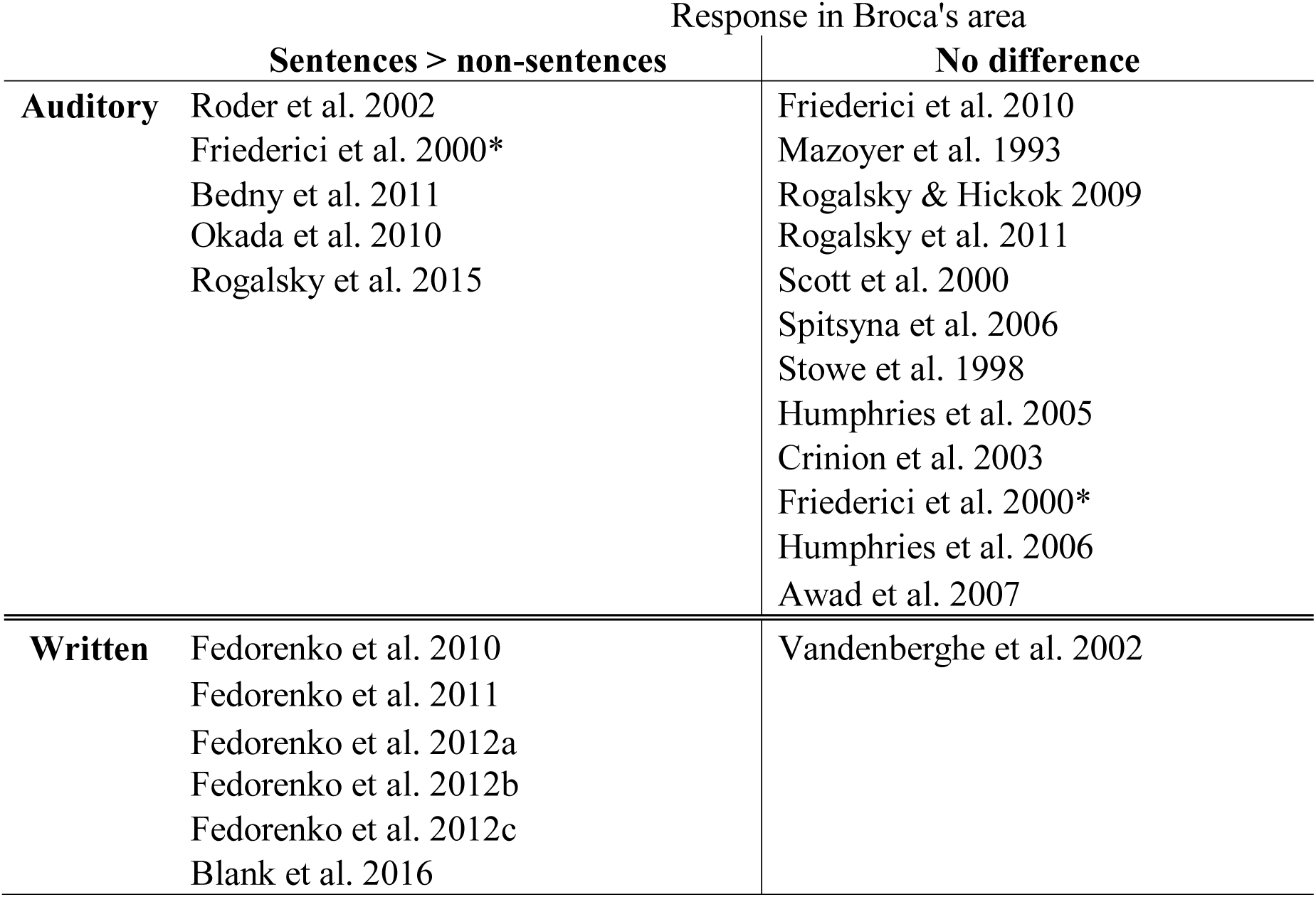
Summary of neuroimaging studies reporting contrasts of sentences versus acoustically-matched control conditions, sorted by stimulus presentation type. Studies included in the “sentences > non-sentences” column report at least one peak activation for this contrast in Broca’s area (i.e. posterior 2/3 of the inferior frontal gyrus). *This study is listed twice because its contrast of normal sentences versus word lists did not activate Broca’s area, but jabberwocky sentences > word lists did activate Broca’s area.

Another potentially important consideration in reconciling functional imaging and lesion results regarding sentence comprehension is functional reorganization. The current study’s participants and those in Dronkers et al.’s (2004) study were chronic stroke patients (i.e. > 6 months and > 1 year post-onset of stroke, respectively) and it can be assumed that their overall language abilities have improved since the initial lesion onset. However, studies of acute stroke patients (i.e. within 48 hours of onset) also identify temporal and inferior parietal regions (approximately Brodmann areas 39 and 40) as the neural substrate for comprehending syntactically complex constructions, such as object-cleft sentences and reversible questions (Magnusdottir et al. 2013; Newhart et al. 2012; Race, Ochfeld, Leigh & Hillis, 2012). Thus, compensatory plasticity does not appear to explain the failure to implicate Broca’s area in sentence comprehension in chronic stroke studies.

### Conclusion

The present study sought to characterize the contributions of damage to Broca’s area and the ATL to agrammatic comprehension. Overall our findings do not support Broca’s area or the ATL being causally involved in processing noncanonical compared to canonical sentences. Specifically, we found:

1. damage to neither Broca’s area nor the ATL resulted in the classic agrammatic comprehension pattern, that is, performance at or near chance level on noncanonical items for both tasks and substantially better on canonical sentence structures.
2. patients with agrammatic comprehension on one or both tasks do not have maximal lesion overlap in Broca’s area or the ATL.
3. (3)Noncanonical sentence comprehension, once variability due to canonical performance is controlled for, implicated posterior temporal-parietal regions, not Broca’s area or the ATL.
4. whole sample VLSM analyses do not implicate Broca’s area in performance on any sentence type in any task, but did implicate Broca’s area in the cognitive task demands of the sentence-picture matching task. The ATL, in combination with posterior temporal-parietal regions, was implicated in performance on both noncanonical and canonical sentences suggesting that the ATL may play an important role in sentence comprehension more broadly.

The present findings also highlight the importance of posterior temporal lobe networks in agrammatic comprehension and sentence comprehension more generally.

## Footnotes

1 It should also be noted that these task-effect studies have almost exclusively only examined stroke patients with aphasia; thus again it is not clear if these patterns would hold in a sample of stroke patients recruited regardless of aphasia diagnosis, and thus would likely include lesions more focal to the areas of interest.

2 The temporal lobectomies were performed on patients with medically intractable epilepsy. These patients have lesions localized in the anterior temporal lobe. As discussed in the introduction, any findings resulting from this population should be interpreted carefully. The results presented here do not change if this group of participants is excluded, but their inclusion allows us to examine (albeit cautiously) the possible involvement of the anterior temporal lobe in sentence comprehension.

## References

Alexander, M.P., Naeser, M.A. & Palumbo, C. (1990). Broca’s area aphasias: aphasia after lesions including the frontal operculum. Neurology, 40, 353–362.

Amunts, K., Schleicher, A., Burgel, U., Mohlberg, H., Uylings, H.B.M., & Zillers, K. (1999). Broca’s region revisited: cytoarchitecture and intersubject variability. Journal of Comparative Neurology, 412, 319–341.

Anwander, A., Tittgemeyer, M., von Cramon, D.Y., Friederici, A.D. & Knosche, T.R. (2007). Connectivity-based parcellation of Broca’s area. Cerebral Cortex, 17, 816–825.

Awad, M., Warren, J.E., Scott, S.K., Turkheimer, F.E. & Wise, R.Jj (2007). A common system for the comprehension and production of narrative speech. Journal of Neuroscience, 27(43), 11455–11464.

Baddeley, A.D., Thomson, N. & Buchanan, M. (1975). Word length and the structure of short-term memory. Journal of Verbal Learning and Verbal Behavior, 14, 575–589.

Baker, E., Blumstein, S.E. & Goodglass, H. (1981). Interaction between phonological and semantic factors in auditory comprehension. Neuropsychologia, 19(1), 1–15.

Barr, D.J., Levy, R., Scheepers, C. & Tily, H.J. (2013). Random effects structure for confirmatory hypothesis testing: keep it maximal. Journal of Memory and Language, 68(3), doi:10.1016/j.jml.2012.11.001

Bates, E., Wilson, S.M., Saygin, A.P., Dick, F., Sereno, M.I., Knight, R.T. & Dronkers, N.F. (2003). Voxel-based lesion-symptom mapping. Nature Neuroscience, 6(5), 448–50.

Bedny, M., Pascual-Leone, A., Dodell-Feder, D., Fedorenko, E. & Saxe, R. (2011). Language processing in the occipital lobe of congentially blind adults. Proceedings of the National Academy of Sciences, 108(11), 4429–4434.

Ben-Shachar, M., Palti, D. & Grodzinsky, Y. (2004). Neural correlates of syntactic movement: converging evidence from two fMRI experiments. NeuroImage, 21(4), 1320–36.

Berndt, R.S., Mitchum, C.C. & Wayland, S. (1997). Patterns of sentence comprehension in aphasia: a consideration of three hypotheses. Brain and Language, 60(2), 197–221.

Berndt, R.S., Mitchum, C.C., & Haendiges, A.N. (1996). Comprehension of reversible sentences in “agrammatism”: a meta-analysis. Cognition, 58, 289–308.

Birembaum, D., Bancroft, L.W. & Felsberg, G.J. (2011). Imaging in acute stroke. Western Journal of Emergency Medicine, 12(1), 67–76.

Blank, I., Balewski, Z., Mahowald, K., Fedorenko, E. (2016). Syntactic processing is distributed across the language system. Neuroimage, 127, 307–323.

Bonner, M.F. & Price, A.R. (2013). Where is the anterior temporal lobe and what does it do? The Journal of Neuroscience, 33(10), 4213–4215.

Bradley, D.C., Garrett, M.E. & Zurif, E.B. (1980). Syntactic deficits in Broca’s aphasia. In: D. Caplan, editor. Biological Studies of Mental Processes. Cambridge: MIT Press.

Brennan, J., Nir, Y., Hasson, U., Malach, R., Heeger, D.J. & Pylkkanen, L. (2012). Syntactic structure building in the anterior temporal lobe during natural story listening. Brain and Language, 120(2), 163–173.

Brennan, J. & Pylkkanen, L. (2012). The time-course and spatial distribution of brain activity associated with sentence processing. Neuroimage, 60(2), 1139–1148.

Brown-Séquard, C. (1876). Dr. Brown-Sequard’s Lectures. Lancet, 2, 203.

Buchsbaum, B.R., Baldo, J., Okada, K., Berman, K.F., Dronkers, N., D’Esposito, M. & Hickok, G. (2011). Conduction aphasia, sensory-motor integration, and phonological short-term memory-an aggregate analysis of lesion and fMRI data. Brain and Language, 119(3), 119–128.

Buchsbaum, B.R. & D’Esposito, M. (2008). The search for the phonological store: from loop to convolution. Journal of Cognitive Neuroscience, 20(5), 765–78.

Buchsbaum, B.R., Ye, D., D’Esposito, M. (2011). Recency effects in the inferior parietal lobe during verbal recognition memory. Frontiers in Human Neuroscience, 5(59).

Caplan, D., Alpert, N. & Waters, G. (1998). Effects of syntactic structure and propositional number on patters of regional cerebral blood flow. Journal of Cognitive Neuroscience, 10, 541–552.

Caplan, D., Alpert, N. & Waters, G. (1999). PET studies of syntactic processing with auditory sentence presentation. Neuroimage, 9, 343–351.

Caplan, D., Alpert, N., Waters, G. & Olivieri, A. (2000). Activation of Broca’s area by syntactic processing under conditions of concurrent articulation. Human Brain Mapping, 9(2), 65–71.

Caplan, D., Chen, E. & Waters, G. (2008). Task-dependent and task-independent neurovascular responses to syntactic processing. Cortex, 44, 257–275.

Caplan, D., DeDe, G. & Michaud, J. (2006). Task-independent and task-specific syntactic deficits in aphasic comprehension. Aphasiology, 20, 893–920.

Caplan, D. & Futter, C. (1986). Assignment of thematic roles by an agrammatic aphasic. Brain and Language, 27, 117–134.

Caplan, D., Michaud, J. & Hufford, R. (2013). Dissociations and associations of performance in syntactic comprehension in aphasia and their implications for the nature of aphasic deficits. Brain and Language 127(1), 21–33.

Caplan, D., Michaud, J., Hufford, R. & Makris, N. (2016). Deficit-lesion correlations in syntactic comprehension in aphasia. Brain and Language, 152, 14–27.

Caplan, D., Stanczak, L. & Waters, G. (2008). Syntactic and thematic constraint effects on blood oxygenation level dependent signal correlates of comprehension of relative clauses. Journal of Cognitive Neuroscience, 20, 643–656.

Caplan, D., Waters, G.S. & Hildebrandt, N. (1997). Determinants of sentence comprehension in aphasic patients in sentence-picture matching tasks. Journal of Speech, Language and Hearing Research, 40(3), 542–555.

Caramazza, A., Capasso, R., Capitani, E. & Miceli, G. (2005). Patterns of comprehension performance in agrammatic Broca’s aphasia: A test of the Trace Deletion Hypothesis. Brain and Language 94(1), 43–53.

Caramazza, A. & Zurif, E.B. (1976). Dissociation of algorithmic and heuristic processes in language comprehension: evidence from aphasia. Brain and Language, 3(4), 572–582.

Crinion, J.T., Lambon-Ralph, M.A., Warburton, E.A., Howard, D. & Wise, R.J. (2003). Temporal lobe regions engaged during normal speech comprehension. Brain, 126(Pt 5), 1193-1201.

Cupples, L. & Inglis, A.L. (1993). When task demands induce “asyntactic” sentence comprehension: a study of sentence interpretation in aphasia. Cognitive Neuropsychology, 10, 201–234.

Damasio, H. (2000). The lesion method in cognitive neuroscience. In: Boller F, Graffman J, Rizzolati G, editors. Handbook of Neuropsychology, 2nd edition. Philadelphia: Elsevier. P 77-102.

Damasio, H. & Damasio, A.R. (1980). The anatomical basis of conduction aphasia. Brain, 103(2), 337–350.

Damasio, H. & Damasio, A.R. (1989). The lesion method in behavioral neurology and neuropsychology. In Feinberg TE, Farah MJ, editors. Behavioral Neurology and Neuropsychology, 2nd edition. New York: McGraw Hill.

Daneman, M. & Newson, M. (1992). Assessing the importance of subvocalization during normal silent reading. Reading and Writing, 4(1), 55–77.

Dapretto, M. & Bookheimer, S.Y. (1999). Form and content: dissociating syntax and semantics in sentence comprehension. Neuron, 24, 292–293.

Drane, D.L., Ojemann, G.A., Ojemann, J.G., Aylward, E., Silbergeld, D.L., Miller, J.W. et al. (2009). Category-specific recognition and naming deficits following resection of a right anterior temporal lobe tumor in a patient with atypical language lateralization. Cortex, 45(5), 630–640.

Dronkers, N.F., Wilkins, D.P., Van Valin, Jr. R.D., Redfern, B.B. & Jaeger, J.J. (2004). Lesion analysis of the brain areas involved in language comprehension. Cognition, 92(1–2), 145–177.

Ellis, A.W., Young, A.W. & Critchley, E.M. (1989). Loss of memory for people following temporal lobe damage. Brain, 112(Pt6), 1469–83.

Fedorenko, E., Behr, M.K. & Kanwisher, N. (2011). Functional specificity for high-level lingustic processing in the human brain. Proceedings of the National Academy of Sciences, 108(39), 16428–33.

Fedorenko, E., Duncan, J. & Kanwisher, N. (2012a). Language-selective and domain-general regions lie side by side within Broca’s area. Current Biology, 22(21), 2059–2062.

Fedorenko, E., Hsieh, P.J., Nieto-Castanon, A., Whitfield-Gabrieli, S. & Kanwisher, N. (2010). New method for fMRI investigations of language: defining ROIs functionally in individual subjects. Journal of Neurophysiology, 104(2), 1177–1194.

Fedorenko, E. & Kanwisher, N. (2011). Some regions within Broca’s area do respond more strongly to sentences than to linguistically degraded stimuli: a comment on Rogalsky and Hickok. Journal of Cognitive Neuroscience, 23(10), 2632–2635.

Fedorenko, E., Nieto-Castanon, A. & Kanwisher, N. (2012b). Lexical and syntactic representations in the brain: an fMRI investigation with multi-voxel pattern analyses. Neuropsychologia, 50(4), 499–513.

Fedorenko, E., Nieto-Castanon, A., Kanwisher, N. (2012c). Syntactic processing in the human brain: what we know, what we don’t know and a suggestion for how to proceed. Brain and Language, 120(2), 187–207.

Fiez, J.A., Damasio, H. & Grabowski, T.J. (2000). Lesion segmentation and manual warping to a reference brain: intra-and interobserver reliability. Human Brain Mapping, 9(4), 192–211.

Frank, R.J., Damasio, H. & Grabowski, T.J. (1997). Brainvox: an interactive, multimodal visualization and analysis system for neuroanatomical imaging. Neuroimage, 5(1), 13–30.

Freud, S. (1891). On Aphasia: a critical study. New York: International Universities Press, Inc.

Fridriksson, J., Bonilha, L. & Rorden, C. (2007). Severe Broca’s aphasia without Broca’s area damage. Behavioral Neurology, 18(4), 237–238.

Fridriksson, J., Fillmore, P., Guo, D. & Rorden, C. (2015). Chronic Broca’s aphasia is caused by damage to Broca’s and Wernicke’s areas. Cerebral Cortex 25(12), 4689–4696.

Friederici, A.D. (2011). The brain basis of language processing: from structure to function. Physiological Reviews, 91(4), 1357–1392.

Friederici, A.D., Kotz, S.A., Scott, S.K. & Obleser, J. (2010). Disentangling syntax and intelligibility in auditory language comprehension. Human Brain Mapping, 31(3), 448–457.

Friederici, A.D., Meyer, M. & von Cramon, D.Y. (2000). Auditory language comprehension: an event-related fMRI study on the processing of syntactic and lexical information. Brain and Language, 75(3), 289–300.

Friederici, A. D., Ruschemeyer, S. A., Hahne, A., & Fiebach, C. J. (2003). The role of left inferior frontal and superior temporal cortex in sentence comprehension: Localizing syntactic and semantic processes. Cerebral Cortex, 13, 170–177.

Frisk, V. & Milner, B. (1990). The relationship of working memory to the immediate recall of stories following unilateral temporal or frontal lobectomy. Neuropsychologia, 28(2), 121– 135.

Glosser, G. & Donofrio, N. (2001). Differences between nouns and verbs after anterior temporal lobectomy. Neuropsychology, 15(1), 39–47.

Goghari, V.M. & MacDonald, A.W. 3rd (2009). The neural basis of cognitive control: response selection and inhibition. Brain and Cognition, 71(2), 72–83.

Goodglass, H. & Kaplan, E. (1983). Boston Diagnostic Aphasia Examination (BDAE). 2nd ed. Philadelphia, PA: Lippincott Williams & Wilkins.

Graff-Radford, J., Jones, D.T., Strand, E.A., Rabinstein, A.A., Duffy, J.R. & Josephs, K.A. (2014). The neuroanatomy of pure apraxia of speech in stroke. Brain and Language, 129, 43–46.

Grodzinsky, Y. (1986). Language deficits and the theory of syntax. Brain and Language, 27, 135–159.

Grodzinsky, Y. (1989). Agrammatic comprehension of relative clauses. Brain and Language, 37(3), 480–499.

Grodzinsky, Y. (1990). Theoretical perspectives on language deficits. Cambridge, MA: MIT Press.

Grodzinsky, Y. (2000). The neurology of syntax: language use without Broca’s area. Behavioral and Brain Sciences, 23(1), 1–21.

Grodzinsky, Y. & Santi, A. (2008). The battle for Broca’s region. Trends in Cognitive Sciences, 12(12), 474–480.

Gutman, R., DeDe, G., Michaud, J., Liu, J. & Caplan, D. (2010). Rasch models of aphasic performance on syntactic comprehension. Cognitive Neuropsychology, 27(3), 230–244.

Harding, I.H., Yucel, M., Harrison, B.J., Pantelis, C. & Breakspar, M. (2015). Effective connectivity within the frontoparietal control network differentiates cognitive control and working memory. Neuroimage, 106, 144–153.

Herrmann, B., Obleser, J., Kalberlah, C., Haynes, J. & Friederici, A.D. (2012). Dissociable neural imprints of perception and grammar in auditory functional imaging. Human Brain Mapping, 33(3), 584–595.

Hertz-Pannier, L., Chiron, C., Jambaque, I., Renaux-Kieffer, V., Van de Moortele, P.F., Delalande, .O. et al. (2002). Late plasticity for language in a child’s non-dominant hemisphere: a pre-and post-surgery fMRI study. Brain, 125(Pt 2), 261-72.

Hickok, G., Buchsbaum, B., Humphries, C. & Muftuler, T. (2003). Auditory-motor interaction revealed by fMRI: speech, music, and working memory in area Spt. Journal of Cognitive Neuroscience, 15(5), 673–82.

Hickok, G., Costanzo, M., Capasso, R. & Miceli, G. (2011). The role of Broca’s area in speech perception: evidence from aphasia revisited. Brain and Language, 119(3), 214–20.

Hickok, G. & Poeppel, D. (2007). The cortical organization of speech processing. Nature Reviews Neuroscience, 8, 393–402.

Holland, R. & Lambon Ralph, M.A. (2010). The anterior temporal lobe semantic hub is a part of the language neural network: selective disruption of irregular past tense verbs by rTMS. Cerebral Cortex, 20(12), 2771–2775.

Huang, C.W., Hayman-Abello, B., Hayman-Abello, S., Derry, P. & McLachlan, R.S. (2014). Subjective Memory Evaluation before and after Temporal Lobe Epilepsy Surgery. PLoS One, 9(4), e93382.

Hughlings Jackson, J. (1866). Notes on the physiology and pathology of language: remarks on those cases of disease of the nervous system in which defect of experession is the most striking symptom. Medical Times and Gazette, i, 659-662.

Humphries, C., Binder, J.R., Medler, D.A. & Liebenthal, E. (2006). Syntactic and semantic modulation of neural activity during auditory sentence comprehension. Journal of Cognitive Neuroscience, 18(4), 665–679.

Humphries, C., Binder, J.R., Medler, D.A. & Liebenthal, E. (2007). Time course of semantic processes during sentence comprehension: an fMRI study. Neuroimage, 36(3), 924–932.

Humphries, C., Love, T., Swinney, D. & Hickok, G. (2005). Response of anterior temporal cortex to syntactic and prosodic manipulations during sentence processing. Human Brain Mapping, 26(2), 128–138.

Jaeger, T. F. (2008). Categorical data analysis: Away from ANOVAs (transformation or not) and towards logit mixed models. Journal of Memory and Language, 59(4), 434–446.

Janecek, J.K., Winstanley, F.S., Sabsevitz, D.S., Raghavan, M., Mueller, W., Binder, J.R. & Swanson, S.J. (2013). Naming Outcome After Left or Right Temporal Lobectomy in Patients with Bilateral Language Representation by Wada Testing. Epilepsy & Behavior, 28(1), 95–98.

Just, M.A., Carpenter, P.A., Keller, T.A., Eddy, W.F. & Thulborn, K.R. (1996). Brain activation modulated by sentence comprehension. Science, 274, 114–116.

Kaan, E. & Swaab, T.Y. (2002). The brain circuitry of syntactic comprehension. Trends in Cognitive Sciences, 6, 350–356.

Kertesz, A. (1982). Western aphasia battery test manual. Psychological Corp.

Kertesz, A. (1997). Recovery of aphasia. In Behavioral Neurology and Neuropsychology, eds. Feinberg, T.E. & Farah, M.J., New York, McGraw Hill, p. 167–182.

Kho, K.H., Indefrey, P., Hagoort, P, van Veelen, C.W., van Rijen, P.C. & Ramsey, N.F. (2008). Unimpaired sentence comprehension after anterior temporal cortex resection. Neuropsychologia, 46(4), 1170–1178.

Lauro, L.J.R., Tettamanti, M., Cappa, S.F. & Papagno, C. (2008). Idiom comprehension: a prefrontal task? Cerebral Cortex, 18(1), 162–170.

Leiken, K. & Pylkkanen, L. (2014). MEG evidence that the LIFG effect of object extraction requires similarity-based interference. Language and Cognitive Processes, 29(3), 381–389.

Linebarger, M.C., Schwartz, M. & Saffran, Ee (1983). Sensitivity to grammatical structure in so-called agrammatic aphasics. Cognition, 13, 361–393.

Love, T. & Oster, E. (2002). On the categorization of aphasic typologies: the SOAP (a test of syntactic complexity). Journal of Psycholinguist Research, 31(5), 503–29.

Magnusdottir, S., Fillmore, P., den Ouden, D.B., Hjaltason, H., Rorden, C., Kjartansson, O. et al. (2013). Damage to left anterior temporal cortex predicts impairment of complex syntactic processing: a lesion-symptom mapping study. Human Brain Mapping, 34(10), 2715–2723.

Marie, P. (1906). The third frontal convolution plays no special role in the function of language. Semaine Medicale, 26, 241–247.

Mazoyer, B.M., Tzourio, N., Frak, V., Syrota, A., Murayama, N., Levrier, O., et al. (1993). The cortical representation of speech. Journal of Cognitivef Neuroscience, 5, 467–479.

Mohr, J.P. (1976). Broca’s area and Broca’s aphasia. In: H. Whitaker & H.A. Whitaker, editors. Studies in Neurolinguistics, volume 1. New York: Academic Press.

Mohr, J.P., Pessin, M.S., Finkelstein, S., Funkenstein, H.H., Duncan, G.W. & Davis, K.R. (1978). Broca’s aphasia: pathological and clinical. Neurology, 28, 311–324.

Newhart, M., Trupe, L.A., Gomez, Y., Cloutman, L., Molitoris, J.J., Davis, C., et al. (2012). Asyntactic Comprehension, Working Memory, and Acute Ischemia in Broca’s Area versus Angular Gyrus. Cortex, 48(10), 1288–1297.

Novick, J.M., Trueswell, J.C. & Thompson-Schill, S.L. (2005). Cognitive control and parsing: reexamining the role of Broca’s area in sentence comprehension. Cognitive, Affective and Behavioral Neuroscience, 5, 263–281.

Ochfeld, E., Newhart, M., Molitoris, J., Leigh, R., Cloutman, L., Davis, C., Crinion, J. & Hillis, A.E. (2010). Ischemia in Broca’s area is associated with Broca’s aphasia more reliably in acute than chronic stroke. Stroke, 41(2), 325–330.

Okada, K., Rong, F., Venezia, J., Matchin, W., Hsieh, I.H., Saberi, K., Serences, J.T. & Hickok, G. (2010). Hierarchical organization of human auditory cortex: evidence from acoustic invariance in the response to intelligible speech. Cerebral Cortex, 20(10), 2486–2495.

Pantasiz, D., Joshi, A., Jiang, J., Shattuck, D.W., Bernstein, L.E., Damasio, H. & Leahy, R.M. (2010). Comparison of landmark-based and automatic methods for cortical surface registration. Neuroimage, 49(3), 2479–93.

Pettigrew, C. & Hillis, A.E. (2014). Role for memory capacity in sentence comprehension: evidence from acute stroke. Aphasiology, 28(10), 1258–1280.

Pillay, S.B., Binder, J.R., Humphries, C., Gross, W.L. & Book, D.S. (2017). Lesion localization of speech comprehension deficits in chronic aphasia. Neurology, 88(10), 970–975.

Race, D.S., Ochfeld, E., Leigh, R. & Hillis, A.E. (2012). Lesion analysis of cortical regions associated with the comprehension of nonreversible and reversible yes/no questions. Neuropsychologia, 50(8), 1946–1953.

Roder, B., Stock, O., Bien, S., Neville, H. & Rosler, F. (2002). Speech processing activates visual cortex in congenitaly blind humans. European Journal of Neuroscience, 16(5), 930–936.

Rogalsky, C., Almeida, D., Sprouse, J. & Hickok, G. (2015). Sentence processing selectivity in Broca’s area: evident for structure but not syntactic movement. Language, Cognition and Neuroscience, 30, 1326–1338.

Rogalsky, C. & Hickok, G. (2009). Selective attention to semantic and syntactic features modulates sentence processing networks in anterior temporal cortex. Cerebral Cortex, 19, 786–796.

Rogalsky, C. & Hickok, G. (2011). The role of Broca’s area in sentence comprehension. Journal of Cognitive Neuroscience, 23(7), 16648–80.

Rogalsky, C., Matchin, W. & Hickok, G. (2008). Broca’s area, sentence comprehension, and working memory: an fMRI study. Frontiers in Human Neuroscience, 2(14), 1–13.

Rogalsky, C., Rong, F., Saberi, K. & Hickok, G. (2011). Functional anatomy of language and music perception: temporal and structural factors investigated using functional magnetic resonance imaging. Journal of Neuroscience, 31(10), 3843–3852.

Rorden, C., Karnath, H.O. & Bonilha, L. (2007). Improving lesion-symptom mapping. Journal of Cognitive Neuroscience, 19(7), 1081–1088.

Santi, A. & Grodzinsky, Y. (2012). Broca’s area and sentence comprehension: a relationship parasitic on dependency, displacement or predictability. Neuropsychologia, 50(5), 821–832.

Santi, A. & Grodzinsky, Y. (2007a). Working memory and syntax interact in Broca’s area. Neuroimage, 37(1), 8–17.

Santi, A. & Grodzinsky, Y. (2007b). Taxing working memory with syntax: bihemispheric modulations. Human Brain Mapping, 11, 1089–1097.

Saykin, A.J., Stafiniak, P., Robinson, L.J., Flannery, K.A., Gur, R.C., O’Connor, M.J. & Sperling, M.R. (1995). Language before and after temporal lobectomy: specificity of acute changes and relation to early risk factors. Epilepsia, 36(11), 1071–1077.

Schwarz, M. & Pauli, E. (2009). Postoperative speech processing in temporal lobe epilepsy: functional relationship between object naming, semantics and phonology. Epilepsy & Behavior, 16(4), 629–633.

Scott, S.K., Blank, C.C., Rosen, S. & Wise, R.J.S. (2000). Identification of a pathway for intelligible speech in the left temporal lobe. Brain, 123, 2400–2406.

Simmons, W.K. & Martin, A. (2009). The anterior temporal lobes and the functional architecture of semantic memory. Journal of the International Neuropsychological Society, 15(5), 645–649.

Slowiaczek, M.L. & Clifton, C. (1980). Subvocalization and reading for meaning. Journal of Verbal Learning and Verbal Behavior, 19, 573–582.

Smith, E.E. & Jonides, J. (1997). Working memory: a view from neuroimaging. Cognitive Psychology, 33, 5–42.

Smith, E.E., Jonides, J., Marshuetz, C. & Koeppe, R.A. (1998). Components of verbal working memory: evidence from neuroimaging. Proceedings of the National Academy of Science, 95, 876–882.

Snijders, T.M., Vosse, T., Kempen, G., Van Berkum, J.J.A., Petersson, K.M. & Hagoort, P. (2009). Retrieval and unification of syntactic structure in sentence comprehension: an fMRI study using word-category ambiguity. Cerebral Cortex, 19, 1493–1503.

Spitsyna, G., Warren, J.E., Scott, S.K., Turkheimer, F.E. & Wise, R.J. (2006). Converging language streams in the human temporal lobe. Journal of Neuroscience, 26, 7328–7336.

Stowe, L.A., Broere, C.A., Paans, A.M., Wijers, A.A., Mulder, G., Vaalburg, W. & Zwarts, F. (1998). Localizing components of a complex task: sentence processing and working memory. Neuroreport, 9(13), 2995–2999.

Stromswold, K., Caplan, D., Alpert, N. & Rauch, S. (1996). Localization of syntactic comprehension by positron emission tomography. Brain and Language, 52, 452–473.

Swanson, S.J., Sabsevitz, D.S., Hammeke, T.A. & Binder, J.R. (2007). Functional magnetic resonance imaging of language in epilepsy. Neuropsychology Review, 17, 491–504.

Swanson, S.J., Binder, J.R., Possing, E.T., Hammeke, T.A., Sabsevitz, D.S., Spanaki, M., et al. (2002). FMRI language laterality during a semantic decision task: age of onset and side of seizure focus effects. Journal of the International Neuropsychological Society, 8, 222.

Swets, J.A. (1964). Signal detection and recognition by human observers, reprint. Los Altos: Peninsula Publishing.

Thothathiri, M., Kimberg, D.Y. & Schwartz, M.F. (2012). The neural basis of reversible sentence comprehension: evidence from voxel-based lesion-symptom mapping in aphasia. Journal of Cognitive Neuroscience, 24(1), 212–222.

Tyler, L.K., Wright, P., Randall, B., Marslen-Wilson, W.D. & Stamatakis, E.A. (2010). Reorganisation of syntactic processing following LH damage: Does RH activity preserve function. Brain, 133, 3396–3408.

Vandenberghe, R., Nobre, A.C. & Price, C.J. (2002). The response of left temporal cortex to sentences. Journal of Cognitive Neuroscience, 14, 550–560.

Venezia, J.H., Saberi, K., Chubb, C. & Hickok, G. (2012). Response bias modulates the speech motor system during syllable discrimination. Frontiers in Psychology, 3, 157.

Wulfeck, B.B. (1988). Grammaticality judgments and sentence comprehension in agrammatic aphasia. Journal of Speech and Hearing Research, 31, 72–81.

Zurif, E.B. (1980). Language mechanisms: a neuropsychological perspective. American Scientist, 68(3), 305–311.

